# Targeting Modulated Vascular Smooth Muscle Cells in Atherosclerosis via FAP-Directed Immunotherapy

**DOI:** 10.1101/2025.03.03.641211

**Authors:** Junedh M. Amrute, In-Hyuk Jung, Tracy Yamawaki, Andrea Bredemeyer, Johanna Diekmann, Sikander Hayat, Wen-Ling Lin, Xianglong Zhang, Xin Luo, Sidrah Maryam, Gyu Seong Heo, Steven Yang, Chang Jie Mick Lee, Caroline Chou, Christoph Kuppe, Kevin D. Cook, Atilla Kovacs, Vishnu Chintalgattu, Danielle Pruitt, Jose Barreda, Nathan O. Stitziel, Paul Cheng, Yongjian Liu, Rafael Kramann, Roger S-Y Foo, Ingrid C. Rulifson, Melissa Thomas, Jixin Cui, Thomas Quertermous, Frank M. Bengel, Simon Jackson, Chi-Ming Li, Brandon Ason, Kory J. Lavine

## Abstract

Vascular smooth muscle cell (VSMC) and immune cell diversification play a central role in driving atherosclerotic coronary artery disease (CAD)^1–3^. However, the molecular mechanisms governing cell state transitions within the neo-intima in human CAD remain poorly understood, and no lipid-independent therapies are currently approved for its treatment. Here, we performed multi-omic single-cell gene expression profiling, epitope mapping, and spatial transcriptomics from 27 human coronary arteries. Our analysis identified fibroblast activation protein (FAP) as a marker of modulated VSMCs within the neo-intima. Genetic lineage tracing in mice confirmed that FAP⁺ cells in the plaque originate from medial VSMCs. Additionally, non-invasive positron emission tomography (PET) imaging in patients with CAD revealed focal FAP uptake in atherosclerotic lesions. Spatial transcriptomics further delineated the distinct localization of VSMC and immune cell subsets within plaques, with FAP⁺ states enriched in the neo-intima. To explore the therapeutic potential of targeting de-differentiated VSMCs, we developed an anti-FAP bispecific T-cell engager (BiTE) and demonstrated that it significantly reduced the plaque burden in multiple mouse models of atherosclerosis. Collectively, our study provides the first single-cell and spatially resolved map of human CAD, establishes FAP as a marker of modulated smooth muscle cells, and demonstrates the broader potential of immunotherapeutics for lipid independent targets in atherosclerotic CAD.

## Introduction

Atherosclerosis, a leading cause of cardiovascular disease-related mortality^4^, is characterized by lipid-rich plaque accumulation, vascular remodeling and inflammation, and smooth muscle cell (SMC) dysfunction^1,5–7^. Medial SMCs maintain vascular integrity but undergo de-differentiation into modulated smooth muscle cells that migrate into the intima and contributing to plaque progression. Targetable mechanisms driving these transitions remain poorly defined. Current coronary artery disease (CAD) treatments focus on lipid-lowering^8–10^, yet many patients still experience disease progression^11–13^. Genome-wide association studies (GWAS) have identified an enrichment of CAD-linked genetic variants associated with SMCs^14,15^, suggesting that targeting SMC state transitions may offer a novel therapeutic approach. However, gaps remain in understanding plaque heterogeneity and modulated smooth muscle cell specification.

Advances in single-cell multiomics and spatial transcriptomics now enable high-resolution profiling of healthy and diseased tissues^16–21^. Prior studies analyzing atherosclerotic plaque nuclei characterized epigenomic and nuclear RNA changes^14,15^ but lacked cellular transcriptomic and surface proteomic data, which are crucial for dissecting lesion heterogeneity. Limited access to fresh coronary tissue has further hindered large-scale single-cell and epitope-based sequencing studies. Single-cell spatial transcriptomics provides an opportunity to map cell types within their anatomical and histopathological context^18,20^. In healthy vessels, the intima, media, and adventitia are distinct^22^; however, during atherosclerosis progression, immune infiltration and stromal cell diversification reshape the media and intimal compartments and drive neointimal growth and plaque instability^1–3,23–26^. Comprehensive single-cell and spatial datasets of human CAD capturing these events are lacking.

Here, we performed Cellular Indexing of Transcriptomes and Epitopes by sequencing (CITE-seq)^27^ and single-cell spatial transcriptomics on healthy and diseased human coronary arteries. We identified fibroblast activation protein (FAP) as a marker of modulated SMCs within the neo-intima, leveraged spatial transcriptomics to elucidate an immune-FAP cell niche, and employed genetic lineage tracing to define their origins. To explore the therapeutic potential of targeting FAP⁺ cells in atherosclerosis, we developed an anti-FAP bispecific T-cell engager (BiTE), that harnesses T-cell cytotoxicity to selectively eliminate FAP-expressing cells^28^. While BiTE technology has been extensively investigated in oncology to direct T cells against tumor-associated antigens^28–30^, its application to cardiovascular disease is unexplored. BiTE treatment resulted in a significant reduction in plaque burden across mouse atherosclerosis models, suggesting that immunotherapeutic depletion of FAP⁺ cells may be a viable strategy for altering disease trajectory. Collectively, this work provides a detailed and comprehensive single-cell and spatially resolved map of human CAD, identifies FAP as a marker of modulated SMCs, and introduces BiTE-based immunotherapy as a potential strategy for targeting atherosclerotic plaques by depleting precise SMC-derived states found in human CAD lesions.

## Results

### Single-cell multi-omic map of human CAD

We performed CITE-seq on 27 human coronary arteries obtained from explanted hearts of patients undergoing heart transplantation (**Fig. 1a, Supplementary Fig. 1a, Supplementary Table 1**). Following rigorous quality control, doublet removal^31^, data integration^32^, and clustering, we identified 150,340 cells spanning 14 distinct cell types (**Fig. 1b, Supplementary Fig. 1b-c**), all of which were present across patient samples (**Supplementary Fig. 1d**). Cell type annotation was achieved using both gene and protein expression profiles (**Fig. 1b-c**). Differential gene expression (DGE) analysis enabled annotation of these populations based on canonical gene and protein markers (**Fig. 1d, Supplementary Tables 2-3**). To identify cell types most relevant to human CAD, we integrated population-level GWAS data^33–35^, overlapping cell-type-specific marker genes with genes predicted to be associated with CAD risk variants. This analysis revealed that stromal populations, including modulated SMCs (modSMCs), endothelial cells, and SMCs/pericytes, exhibit the highest enrichment for CAD-associated genetic risk (**Fig. 1d**). Notably, the SMC compartment displayed the greatest enrichment for disease-linked genes, reinforcing prior findings^14,15^ that highlight its central role in driving CAD pathogenesis.

**Figure 1.**
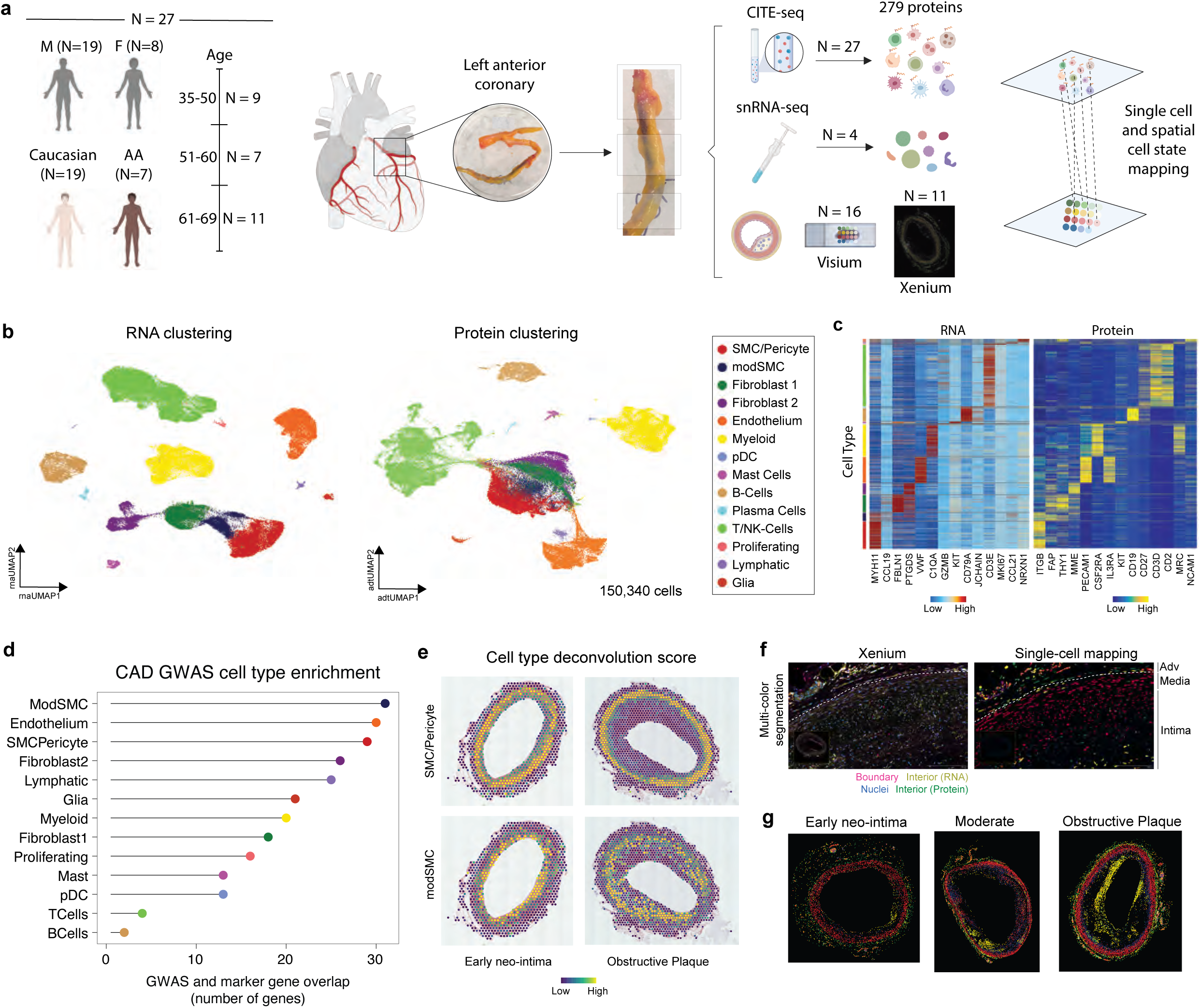
Single-Cell and Spatial Multi-Omics Analysis of Human Coronary Arteries in Health and Disease. (a) Study design using human coronary arteries from explanted hearts for single cell isolation and CITE-seq, histopathology and Visium FFPE transcriptomics, and Xenium spatial transcriptomics. (b) UMAP plot using gene (left) and protein (right) expression. (c) Heatmap of differentially expressed genes and proteins. (d) Overlap of genes linked to CAD GWAS disease variants and marker genes for cell types. (e) Tangram voxel deconvolution scores for SMC/Pericyte and modSMC in a mildly diseased and obstructed vessel. (f) Multi-color cell segmentation in a diseased coronary artery colored by stain (left) and segmented cells mapped onto the CITE-seq reference (right) colored by imputed annotation in Fig. 1b. (g) Example of mild, moderate, and severe plaque Xenium sections colored by reference mapped cell type annotations. For differential gene and protein expression, the Wilcoxon Rank Sum test is used to adjust the p-value.

### Spatial map of human CAD

To characterize the spatial landscape of human CAD, we performed spatial transcriptomics on healthy vessels with early neo-intima (n=4), moderate lesions (n=8), and ruptured or obstructed lesions (n=4) using the Visium platform on formalin-fixed paraffin-embedded (FFPE) tissue (**Supplementary Fig. 2a-c**). Spatial deconvolution was carried out using tangram^36^, leveraging our CITE-seq map as a reference across lesion types (**Supplementary Fig. 2d**). This analysis revealed enriched SMC/pericyte mapping scores in the medial layer and an expansion of modulated SMCs (modSMCs), myeloid cells, and T/NK cells within the neo-intima (**Fig. 1e, Supplementary Fig. 2d**). In healthy vessels, immune cells and fibroblasts were enriched in the adventitia, with early modSMC expansion observed in the developing neo-intima (**Supplementary Fig. 2d**). In more advanced lesions, including moderate and severely obstructed plaques, we detected a pronounced accumulation of modSMCs, myeloid cells, and T/NK cells within the neo-intima and necrotic core (**Supplementary Fig. 2d**). While the Visium platform enables transcriptome-wide characterization of tissue, its spatial resolution is limited to 55 µm voxels, which is too large for single-cell mapping. To overcome this limitation, we designed a custom Xenium spatial transcriptomics panel incorporating 100 genes identified from our CITE-seq dataset to differentiate cell types (**Fig. 1c**) alongside the base panel (**Supplementary Table 4**), and applied it to human CAD vessels (n=11) (**Supplementary Fig. 3a**). Multi-color imaging with our gene panel and cell segmentation probes (interior RNA, interior protein, boundary, nuclear staining) allowed us to precisely map cell types identified from CITE-seq reference map into space (**Fig. 1f-g**, **Supplementary Fig. 3b**). This approach facilitated the mapping of immune and stromal cell populations across lesion groups (**Fig. 1f-g**, **Supplementary Fig. 3b-d**) and revealed a marked expansion of myeloid cells within the neo-intima of the most severe lesions (**Supplementary Fig. 3c-d**).

### FAP enriched on modulated SMCs

We performed unbiased clustering to identify 14 transcriptionally distinct stromal cell states (**Fig. 2a**), each defined by unique gene and protein signatures (**Fig. 2b, Supplementary Fig. 4a, Supplementary Tables 5-6**). These cell states included pericytes (*RGS5, STEAP4, CD36*), four SMC subtypes (SMC1: *RERGL, PHLDA2, BCAM*; SMC2; *MYH11, CNN1, PLN*; SMC3: *ACTG2, MYH10, ITGA8*; SMC4: *IFI44L, IFIT3, MX1*), and 2 modulated SMC subtypes (fibromyocytes/FMCs: *IGFBP2, VCAN, TNFRSF11B*; chondromyocytes/CMCs: *CCL19, COMP, POSTN*). Additionally, we identified seven fibroblast subtypes (Fib1: *PLA2G2A, MFAP5, PCOLCE2*; Fib2: *CFD, C7, FBLN1*; Fib3: *VWF, EGFL7, ACKR1*; Fib4: *CXCL14, LEPR, RARRES2*; Fib5: *ANGPTL7, APOD, PTGDS*; Fib6: *SLC2A1, KRT19, ITGA6*; Fib7: *ABCA8, ABCA10, CDH19*) (**Supplementary Fig. 4a, Supplementary Tables 5-6**). Broadly, SMC1-3 exhibited an enrichment of contractile markers such as *MYH11* and *ACTA2*^1,14,37^, whereas FMCs and CMCs were characterized by synthetic and fibrotic gene signatures^1,38,39^ such as *TNFRSF11B, MGP,* and *POSTN* (**Fig. 2b, Supplementary Fig. 4a**). From our Visium and Xenium spatial transcriptomics data we see that SMC1-3 are enriched within the media, FMC/CMC within the intima, and fibroblasts within the adventitia (**Supplementary Fig. 2-3**). CAD GWAS enrichment analysis revealed the strongest association with FMC marker genes, consistent with recent findings showing an enrichment of CAD-associated variants within FMC-specific enhancers^14^. Notably, the FMC and CMC cell states exhibited high expression of the surface protein FAP (**Fig. 2d**). Immunofluorescence analysis of human atherosclerotic lesions confirmed FAP protein expression, with strong enrichment in the intimal layer and no detectable expression in the media (**Fig. 2e**). Additionally, rank-sorted GTEx bulk tissue expression^40^ (log_10_(TPM + 1)) demonstrated robust FAP expression across multiple arterial beds, including the aorta, coronary artery, and tibial artery tissue (**Fig. 2f**).

**Figure 2.**
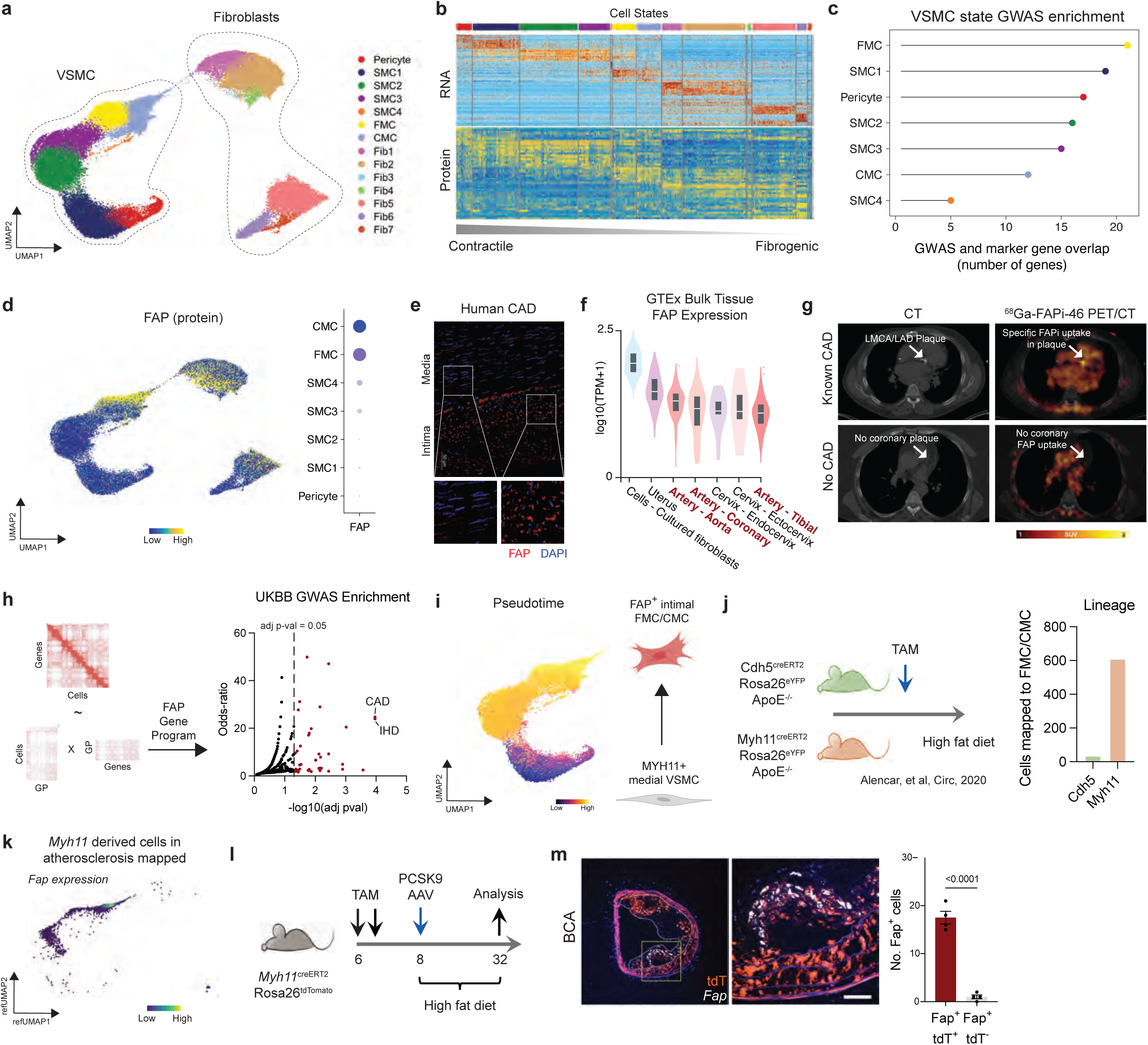
FAP-Associated Stromal Cell States in Atherosclerosis. (a) UMAP plot colored by stromal cell states. (b) Heatmap of differentially expressed genes and proteins across stroma cell states. (c) Overlap of genes linked to CAD GWAS disease variants and marker genes for stromal cell states. (d) FAP protein expression from CITE-seq panel (left) and dotplot of FAP protein expression by cell state (right). (e) Immunofluorescence staining for FAP with DAPI nuclei staining in a diseased human coronary artery; zoom-in on a medial and intimal segment. (f) *FAP* expression from GTEx bulk tissue data ranked by log_10_(TMP + 1). (g) Top row: Representative example of a patient with known coronary artery disease and chronic ischemic cardiomyopathy. CT imaging shows a sclerotic LMCA / LAD plaque. Molecular PET-imaging using ^68^Ga-FAPi-46 reveals specific FAPi-uptake in the calcified plaque (matching CT finging, both indicated by blue arrows). Bottom row: Representative example of a patient without known CAD. No coronary plaques were detected in CT. ^68^Ga-FAPi-46 PET/CT shows no specific tracer uptake in coronary arteries. Blue arrows indicate the LAD. (h) UK biobank GWAS enrichment of genes within the *FAP* cell state (FMC/CMC); the odds ratio is derived from running Fisher exact test for random gene sets to compute a mean rank and standard deviation from the expected rank for each term in the gene-set library and calculating a z-score to assess the deviation from the expected rank, the adjusted p-value is computed using the Bejamini-Hochberg method for correcting for multiple hypotheses testing. (i) Palantir pseudotime overlaid on the UMAP plot for Pericyte, SMC1-4, FMC, and CMC states. (j) *Cdh5* and *Myh11* lineage positive cells from published murine atherosclerosis model mapped onto human FMC/CMC states with quantification of number of cells which map to the FMC/CMC state. (k) *Fap* expression density plot in mouse data reference mapped onto human stromal UMAP highlighting expression in the FMC/CMC compartment. (l) Study schematic for in vivo lineage tracing of *Myh11*^+^ SMCs in a murine atherosclerosis model. (m) RNA in situ hybridization for *Fap* and tdTomato staining in the brachiocephalic artery plaque lesion cross-section after 24 weeks of HFD feeding. Outlined area indicate the region magnified in the next panel (left). Quantification of the number of Fap^+^ cells in plaque lesion, n=4/group (right). Scale bars, 50 µm in Fig. 5m; data were analyzed with unpaired nonparametric Mann-Whitney test. For differential gene and protein expression, the Wilcoxon Rank Sum test was used to adjust the p-value.

### FAPI PET/CT in Human Coronary Arteries

To investigate the translational relevance of FAP^+^ SMC states, we re-analyzed FAPi-targeted PET/CT imaging data from patients with coronary artery disease using the specific ligand 68Ga-FAPI-46^41^. Four coronary territories (LMCA, LAD, LCX and RCA) were visually assessed for presence of calcified plaques or stents in 15 patients (60 coronary arteries analyzed in total). 58% (35/60) of coronary arteries presented with plaques or stentings. 40% of the identified affected vessels showed elevated focal 68Ga-FAPi-46 uptake matching the respective plaque or stent (**Fig. 2g**). These findings highlight potential clinical use of a FAP tracer to identify modulated SMCs within coronary lesions.

### FAP associated gene network

To identify genes enriched in the FAP^+^ cell state, we conducted a transcriptome-wide correlation analysis to determine genes whose expression correlates with FAP protein levels at the single-cell level within pericytes, SMC1-3, and FMC/CMC states. We rank-ordered the top 30 genes that showed a significant correlation (Pearson coefficient > 0.3, adjusted p-value < 0.05) with FAP protein expression (**Supplementary Fig. 4b, Supplementary Table 7**). Among these, *MGP, LUM, VCAN, F2R, OMD, LTBP2*, and *FAP* itself emerged as top candidates (**Supplementary Fig. 4b-c, Supplementary Table 7**). Several of these genes play key roles in extracellular matrix remodeling and fibrosis – *MGP* has been implicated in vascular calcification^42^, while *VCAN* and *LUM* are critical for maintaining extracellular matrix integrity^43,44^. *LTBP2* is involved in TGF-β signaling^45^, further highlighting the potential pathological significance of FAP^+^ cell states. Additionally, we identified five surface proteins that were significantly correlated with FAP protein levels (Pearson coefficient > 0.3, adjusted p-value < 0.05): ITGA2, CD200, THY1, CD63, and F3 (**Supplementary Fig. 4d**). Notably, ITGA2 protein expression was highly enriched in FMC/CMC states (**Supplementary Fig. 4e**) and prior work has implicated ITGA2 as a key protein in mediating cell adhesion and ECM remodeling^46^. Gene Ontology analysis of the FAP gene program revealed enrichment in pathways related to phosphatidylinositol 3-kinase (PI3K) signaling and extracellular matrix reorganization (**Supplementary Fig. 4f, Supplementary Table 8**). The PI3K pathway is a central regulator of cell survival, proliferation, and migration, processes that are critically involved in vascular remodeling and plaque progression^47,48^. Dysregulation of this pathway has been implicated in atherosclerosis, supporting the idea that FAP^+^ cells may contribute to disease pathogenesis through enhanced matrix remodeling and maladaptive cellular responses. Moreover, GWAS enrichment analysis using UK Biobank data showed a significant association between FAP^+^ cell states (FMC/CMC) and coronary artery disease (CAD) and ischemic heart disease (**Fig. 2h, Supplementary Table 9**).

### FAP^+^ cells are derived from medial vascular SMCs

To infer potential vascular SMC lineage trajectories in human CAD we used Palantir^49^. This analysis indicated that contractile SMCs (SMC1-3) have low pseudotime and high entropy values suggesting that they may represent a plastic population. In contrast, FMC and CMC cell states displayed high pseudotime values suggesting that they may be terminal cell states (**Fig. 2i**). We found that the contractile genes *MYH11* and *ACTA2* decreased, while synthetic genes within the FAP gene network (**Supplementary Fig. 4b**) including *VCAN, LTBP2, TNFRSF11B,* and *MGP* increased along pseudotime (**Supplementary Fig. 5a**) implying that modulated SMCs differentiated from vascular SMCs. We used two orthogonal methods to validate these informatic predictions. First, we leveraged published single cell RNA sequencing datasets that included lineage tracing^50^ of SMCs (*Myh11^creERT2^Rosa26^eYFP^*) and endothelial cells (*Cdh5^creERT2^Rosa26^eYFP^*) in a murine atherosclerosis model. We re-processed and mapped these data onto our human CAD CITE-seq map to annotate cell types (**Supplementary Fig. 5b-d**). We observed strong mapping for Pericytes/SMCs, endothelial, and myeloid populations and poor mapping of adventitial fibroblasts (**Supplementary Fig. 5d**). These findings are consistent with prior work^17^ and highlights a discrepancy between human and mouse perivascular fibroblasts. Mapping of vascular SMC and endothelial cell lineages revealed that FAP^+^ modulated SMCs (FMCs/CMCs) were derived from medial vascular SMCs (**Fig. 2j, Supplementary Fig. 5e-f**). *Fap* expression was selectively observed within the FMC/CMC subset of vascular SMC-derived cells in murine atherosclerosis (**Fig. 2k**). Independently, we lineage traced medial vascular SMCs using *Myh11^creERT2^Rosa26^tdTomato^* mice that were later injected with AAV8-D377Y-mPCSK9 and fed a high-fat diet (HFD) for 24 weeks to induce atherosclerosis (**Fig. 2l**). We then performed *Fap* RNAscope in the brachiocephalic artery and found that most *Fap*^+^ cells in the plaque are tdTomato^+^ (derived from *Myh11* lineage) (**Fig. 2m**). Additionally, we reanalyzed published single-cell lineage tracing data^37^ from *Myh11*^creERT2^Rosa26^tdTomato^ in a murine atherosclerosis model (**Supplementary Fig. 5g**), selecting tdTomato^+^ lineage-traced cells. Notably, we observed increased Fap expression after 16 weeks of HFD feeding in these cells (**Supplementary Fig. 5h**). To validate these findings, we performed *Fap* RNAscope and COL1A1 (for staining collagen) immunofluorescence staining in the aortic root of 16 weeks of HFD fed *Apoe^-/-^* mice, confirming elevated *Fap* expression within atherosclerotic plaque lesions (**Supplementary Fig. 5i**). Collectively, our results show that *Fap*^+^ cells in atherosclerosis are derived from medial vascular SMCs.

### Distinct spatial niches in CAD

To dissect spatially restricted niches in human CAD lesions, we performed Visium FFPE spatial transcriptomics on both healthy and diseased vessels (**Supplementary Fig. 2**). We generated consensus meta-programs (MPs) that were conserved across samples using non-negative matrix factorization (NMF) to decompose the data into gene programs^51^ and multiNMF to account for batch effects across samples and combine gene programs into MPs (**Supplementary Fig. 6a, Supplementary Table 10**). We represented each spatial voxel by its MP score and used these components to generate a UMAP visualization, with nearest-neighbor clustering, to identify conserved spatial niches (**Fig. 3a, Supplementary Fig. 6b-c**). Differential gene expression analysis revealed key marker genes associated with each spatial niche (**Supplementary Fig. 6b-c**). Furthermore, we observed that the MP signatures corresponded to different cell populations, as determined by tangram deconvolution (**Supplementary Fig. 6d**). Specifically, MPs 3-6 and 10 were enriched in SMC/Pericytes, MPs 3-5 in modulated SMCs, MP7 in myeloid cells, MPs 1 and 8 in endothelial cells, and MPs 2 and 9 in fibroblasts (**Supplementary Fig. 6d**).

**Figure 3.**
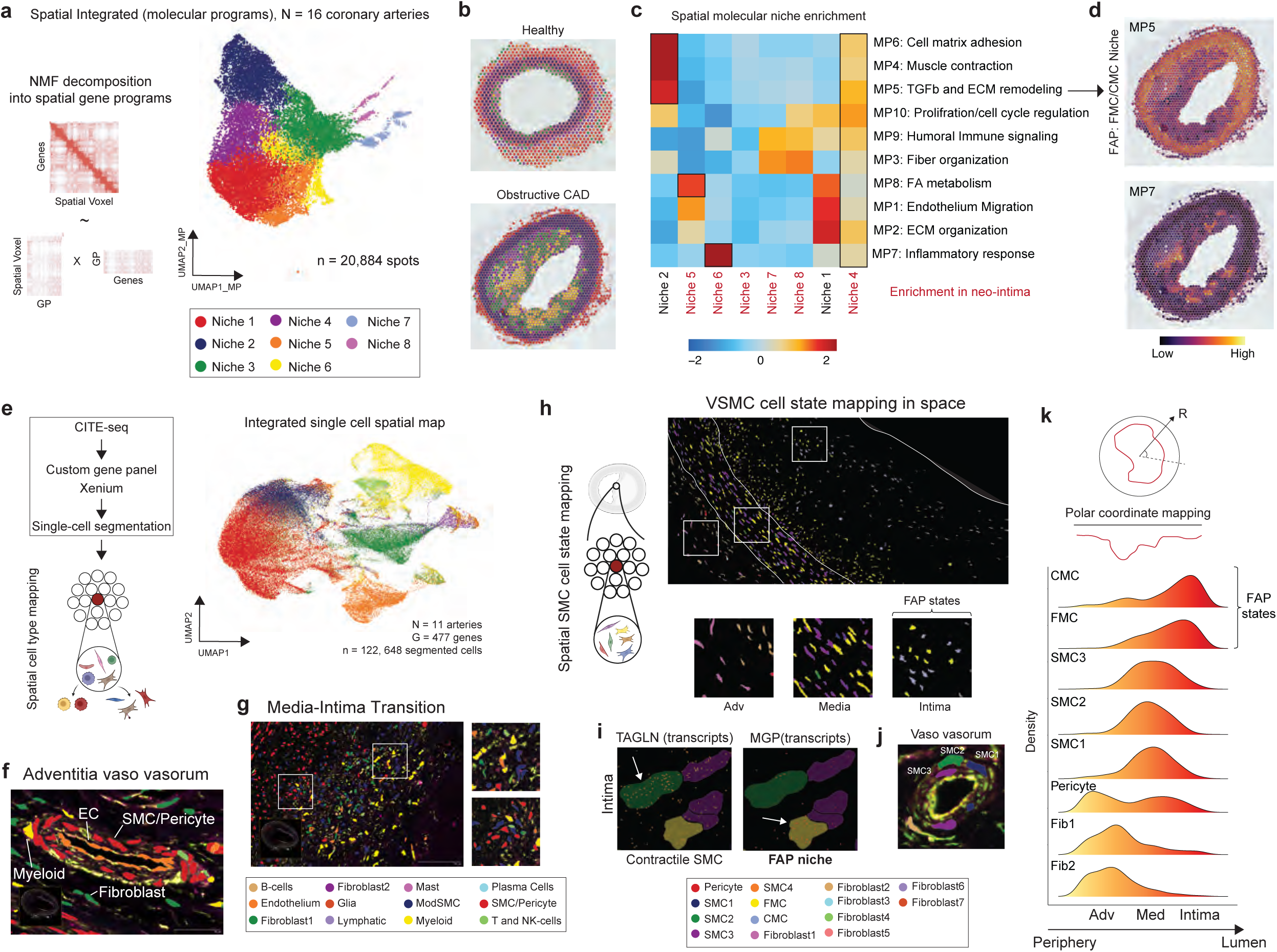
Spatial Niches and Stromal Cell Dynamics in Healthy and Diseased Coronary Arteries. (a) NMF decomposition of spatial voxels (left) and UMAP plot colored by conserved spatial niches from consensus MPs integrated across n=16 samples. (b) Example of a healthy and diseased CAD with spatial voxels colored by Niches from (a). (c) Heatmap of MP gene set scores across Niches from (a) with associated pathways annotated. (d) MP5 and MP7 gene set score in diseased CAD sections. (e) Schematic of analysis framework for Xenium data (left) and UMAP plot of integrated Xenium single segmented cells from n=11 samples colored by reference mapped annotations from Fig. 1b (right). (f) Zoom in on the vaso vasorum in the adventitia colored by cell type. (g) Neo-intima microenvironment colored by cellular stains and cell type annotations. (h) Reference mapping of spatial SMC/Pericytes, modSMCs, and Fibroblasts onto CITE-seq stromal cell state reference (Fig. 2a) annotated in space in a diseased vessel with a zoom in on the adventitia (enriched with fibroblasts), media (contractile SMC, FMC), and neo-intima (FMC/CMC). (i) *TAGLN* and *MGP* transcripts in a zoom in of stromal cells in proximity within the neo-intima. (j) Zoom in on the vaso vasorum in the adventitia colored by SMC cell states. (k) Lumina-to-periphery deconvolution with quantification of stromal cell density along distance from the center of the lumen. For differential gene and protein expression, the Wilcoxon Rank Sum test is used to adjust the p-value.

To assign MPs to their corresponding spatial niches and anatomic location, we first co-registered annotated special niches to histopathology images and found that niche 1 was enriched in the adventitia, niche 2 in the media, and niches 3-8 in the neo-intima (**Fig. 3b, Supplementary Fig. 6e**). Next, to link spatial niches to MPs, we performed gene set enrichment analysis. This revealed enrichment of MPs in niche 1: MPs 1, 2, and 8; niche 2: MPs 4-6; niche 4: MPs 1-10, niche 5: MP8; niche 6: MP7; and niche 7-8: MPs 3 and 9. Of particular interest, was MP5 (associated with TGF-β and ECM remodeling, enriched in niches 2 and 4), which corresponded with FAP^+^ modulated SMC states (FMCs/CMCs). MP5 enriched spatially niches were juxtaposed to MP7 enirched spatial niches (associated with inflammatory response, enriched in niches 4 and 6) at the media-intima boundary (**Fig. 3c-d**). Additionally, PROGENy pathway enrichment^52^ highlighted NF-κB signaling as significantly enriched in the neo-intima and absent in healthy vessels (**Supplementary Fig. 6f**). Notably, the FAP gene module was also enriched in diseased arteries and absent from healthy vessels (**Supplementary Fig. 6g**). These findings provide valuable insights into the spatial organization of CAD lesions, identifying specific molecular signatures present in the atherosclerotic microenvironment.

### Modulated SMCs occupy distinct spatial niches

While Visium empowers transcriptome-wide spatial profiling, it lacks single-cell resolution and thus limits the precise identification of individual cell states in dense microenvironments such as the atherosclerotic plaque. To overcome this limitation and dissect the spatial landscape of CAD at single-cell resolution, we employed Xenium with a custom gene panel (**Supplementary Table 4**) derived from our CITE-seq atlas. After cell segmentation, we assigned transcripts to individual cells and assigned annotations based on mapping to our CITE-seq reference (**Fig. 3e**). Xenium enables high-resolution identification of key cell types within their microenviroment (**Fig. 1f-g, Supplementary Fig. 3b-d**). In the adventitia, we identified distinct populations of macrophages, endothelial cells, fibroblasts as well as SMCs and pericytes that form the vaso vasorum (**Fig. 3f**). In the media and neo-intima, we revealed a high degree of cellular heterogeneity with close approximation of SMCs, modulated SMCs, myeloid cells, and T/NK-cells (**Fig. 3g**). We chose to focus on SMCs, pericytes, modulated SMC, and fibroblast populations to explore their spatial localization (**Supplementary Fig. 7a**). We observed enrichment of fibroblasts in the adventitia, SMC3 in the media, FMCs in the media and neo-intima, and CMCs in the neo-intima (**Fig. 3h, Supplementary Fig. 7b**). Additionally, we visualized higher transcript counts for *TAGLN* in contractile SMC states (SMC 2/3) and greater *MGP* transcript counts in FMCs (**Fig. 3i**). Key genes associated with the FAP module, including *MGP, VCAN*, and *LTBP2*, were enriched in the neo-intima and showed weaker expression in healthy vessels (**Supplementary Fig. 7c**). In the adventitial vasa vasorum, we were able to delineate distinct SMC cell states (SMC 1-3) in adjacent layers of the vessel wall, further illustrating the power of Xenium in characterizing single-cell states in spatial context (**Fig. 3j**). To quantify the spatial orientation of stromal cell states relative to the vessel lumen, we defined the lumen center, mapped the cells using polar coordinates, and calculated the distance from the lumen center (**Fig. 3k**). We found that CMCs, FMCs, SMCs 1-3, Pericytes, and Fibroblasts 1-2 were spatially organized from lumen/intima to adventitia, respectively (**Fig. 3k**). With the Xenium data we were able to resolve spatially organized cellular hierarchies within concentrated microenvironments.

### FAP^+^ cells and macrophages co-localize in plaques

Single-cell spatial transcriptomics provides a unique opportunity to characterize cellular co-localization within human tissue. To explore cell-cell interactions in atherosclerotic lesions, we employed MISTy^53^ predictive modeling. Among the myeloid cells, Mac4 strongly co-localized with FMCs (**Fig. 4a**) while CMCs co-localized with classical monocytes, Mac1, and Mac6. By integrating data across samples, conducting unbiased clustering, and performing differential expression analysis, we identified 12 transcriptionally distinct myeloid subsets (**Fig. 4b, Supplementary Fig. 8a, Supplementary Table 12**). Surface protein analysis revealed that LILRA4 is enriched on tissue-resident macrophages, while CCR2 is predominantly expressed on monocytes, inflammatory macrophages (Mac1), and dendritic cells (**Supplementary Fig. 8b, Supplementary Table 13**). Notably, the Mac4 subset is enriched in genes involved in lipid metabolism, including *FABP4, FABP5, GPNMB, CTSL*, and *SPP1* (**Fig. 4c**). At the protein level, CD276 is prominently expressed on Mac4 cells (**Supplementary Fig. 8b**). We also identified CAD-enriched genes from GWAS data^33,35^ expressed in macrophages, notably *LIPA*, which is enriched in Mac4 and expressed in the neo-intima (**Supplementary Fig. 8c-d**). Single-cell segmentation within the neo-intima showed enriched *GPNMB* transcripts in Mac4 cells and *MGP* transcripts in FMCs (**Fig. 4d-e**). A strong co-localization between the FAP module and the Mac4 gene signature was also observed from the Visium FFPE data (**Supplementary Fig. 8e**) bolstering the conclusion that Mac4 and FMCs co-localize in CAD lesions. To further characterize myeloid cell states in space, we mapped segmented macrophages from the Xenium data onto our CITE-seq myeloid cell state map (**Fig. 4b**). We observed that Mac4 is located in the neo-intima (**Fig. 4f**). As an orthogonal approach, we used a spatial neo-intimal gene signature derived from CAD vessels, which was absent in healthy arteries, and found that this signature predominantly mapped to the Mac4 subset (**Fig. 4g**). Gene ontology biological process enrichment analysis of Mac4 marker genes revealed pathways related to LDL handling and fatty acid metabolism (**Fig. 4h**, **Supplementary Table 14**). Interestingly, the Mac4 gene signature was dynamically modulated in an in vivo murine model of atherosclerosis^54^, with increased expression following HFD feeding and a reduction upon ApoB injection (**Fig. 4i**).

**Figure 4.**
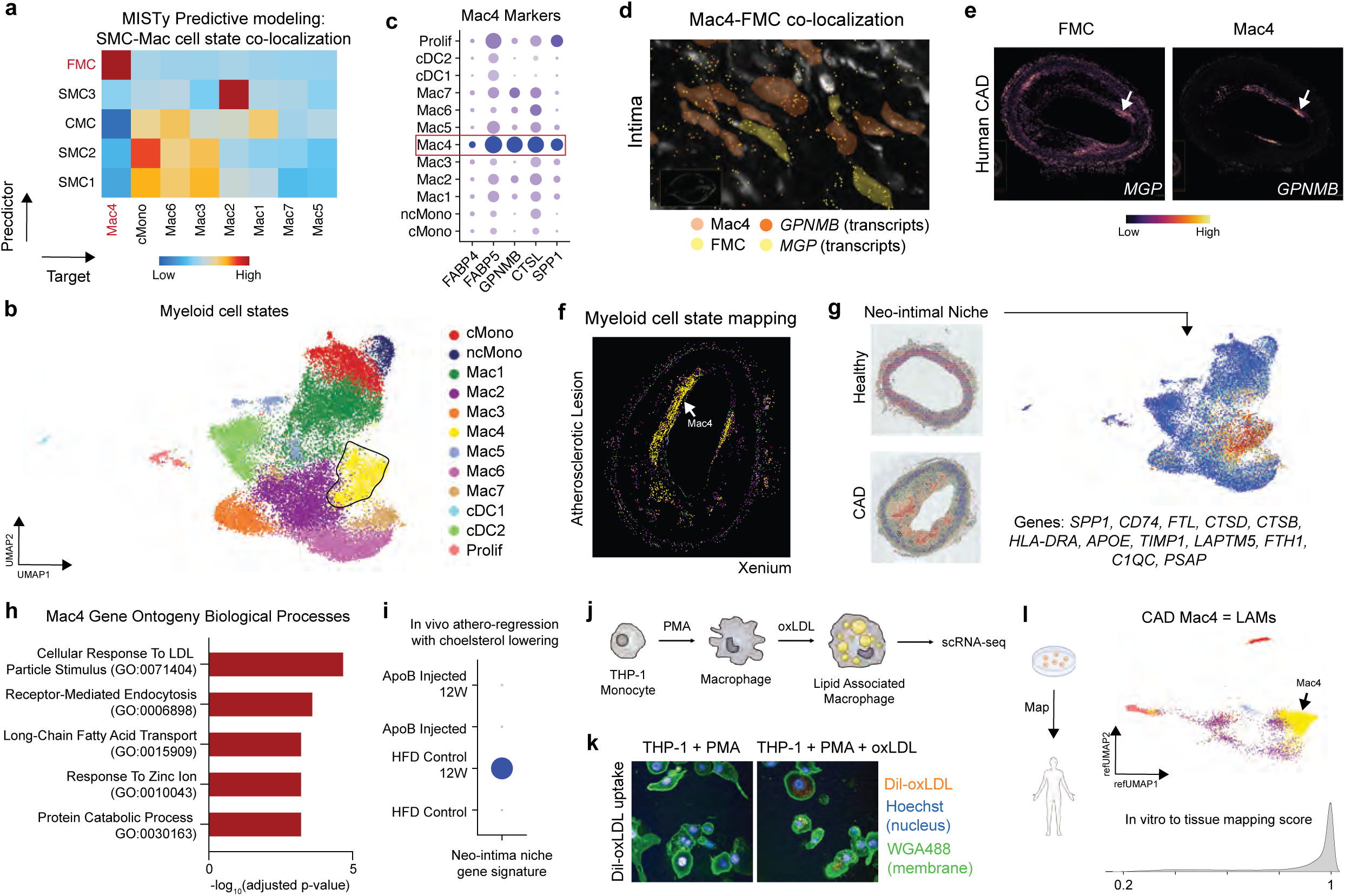
FAP Niche and Myeloid Cell Interactions in Atherosclerosis. (a) Cell state MISTy predictive analysis (SMC1-4, FMC, CMC and Myeloid cell states; atherosclerotic vessels) heatmap of predictor cell (*y* axis) and target cell (*x* axis) from Xenium data. (b) UMAP plot of myeloid cell states. (c) DotPlot of genes enriched within Mac4. (d) Zoom in within the neo-intima showing FMC and Mac4 co-localization with *MGP* and *GPNMB* transcripts. (e) Spatial density plot of *MGP* and *GPNMB* in an atherosclerotic human CAD vessel. (f) Myeloid cell states mapped into space with Xenium colored by annotations in Fig. 4(b). (g) Gene set score for neo-intimal niche in a healthy and diseased vessel (left) and overlaid on myeloid UMAP plot. (h) Gene Ontology biological processes pathway enrichment for Mac4 marker genes; the adjusted p-value is computed using the Bejamini-Hochberg method for correcting for multiple hypotheses testing. (i) DotPlot of gene signature from Fig. 4(g) in published myeloid single cell data from a murine atherosclerosis regression model. (j) Schematic for generation of LAMs in vitro. (k) Immunofluorescence plot of dil-oxLDL uptake in THP-1 + PMA and THP-1 + PMA + oxLDL treated cells. (l) Reference mapping of in vitro LAMs onto myeloid cell sates from Fig. 4(b) (top) with ridge plot of the mapping score (bottom). For differential gene and protein expression, the Wilcoxon Rank Sum test is used to adjust the p-value.

### CAD intimal macrophages resemble lipid associated macrophages

We then compared the transcriptional signature of Mac4 cells localized within human CAD plaques to an established model of foam cells. THP-1 cells were treated with PMA followed by oxLDL (**Fig. 4j-k, Supplementary Fig. 9a-b**) and single cell RNA-sequencing was performed. After quality control, dimensional reduction, data integration, and cell clustering we found 7 distinct cell states of in vitro macrophages (IVMac) 1-7 (**Supplementary Fig. 9d-e, Supplementary Table 15**) These included IVMac1 (*FABP4, GPNMB, NPTX1*), IVMac2 (*TYMS, MYBL2, FBXO5*), IVMac3 (*STARD4, MSMO1, DERL3*), IVMac4 (*PTTG1, PRTN3, CDC20*), IVMac5 (*TOP2A, CDK1, BIRC5*), IVMac6 (*MX1, ISG15, OAS2*), and IVMac7 (*MIF, H2AFZ, STMN1*). Notably IVMac1 and IVMac3 expanded after dil-oxLDL and oxLDL treatment (**Supplementary Fig. 9f**). We plotted the Mac4 gene signature in the in vitro single cell data and found enrichment in IVMac1 and IVMac3 states (**Supplementary Fig. 9g**). Furthermore, we reference mapped the in vitro oxLDL treated macrophages onto the human myeloid cells from human CAD and observed marked enrichment within Mac4 with a strong mapping score. These data indicate that the Mac4 cluster present in human plaques resemble lipid associated macrophages and that established in vitro models appear to be an appropriate surrogate to study this population (**Fig. 4l**).

### Anti-FAP BiTE treatment reduces atherosclerotic plaque burden in vivo

To assess the therapeutic potential of targeting *Fap*^+^ cells, we designed an anti-FAP bispecific T-cell engager (BiTE) antibody that incorporates an anti-FAP monoclonal antibody binder and an anti-CD3ɛ monoclonal antibody that engages and activates T-cells (**Fig. 5a**). We first tested this therapy in an acute murine model^55^ of atherosclerosis by using *Apoe^-/-^*mice subjected to HFD feeding and Angiotensin II infusion via osmotic mini-pumps, with weekly administration of either control or anti-FAP BiTE (**Fig. 5b**). After four weeks of treatment, we observed no significant change in plasma cholesterol levels between experimental groups (**Fig. 5c**). However, anti-FAP BiTE treated mice showed a significant reduction in plaque burden within the aortic arch quantified by enface aortic dissection (**Fig. 5d**). To further evaluate the efficacy of anti-FAP BiTE therapy in a chronic state of disease set, we extended the HFD feeding for 16 weeks. Beginning at week 12, mice were treated with weekly administration of either control or anti-FAP BiTE for four weeks (**Fig. 5e**). Consistent with the acute model, plasma cholesterol levels remained unchanged between group (**Fig. 5f**). When compared with controls, treatment of anti-FAP BiTE also resulted in significant decrease in atherosclerotic plaque burden in the aortic arch by enface aortic dissection (**Fig. 5g**). After the end of the treatment period, Fap expression was assessed by quantifying number of *Fap*^+^ cells in the fibrous cap normalized to atherosclerotic plaque size. We found that anti-FAP BiTE-treated animals showed a significant reduction in the number of *Fap*^+^ cells within aortic root plaques compared to controls (**Fig. 5h-i)**. To assess relative plaque stability, we quantified the percent ACTA2^+^ area in fibrous cap normalized to atherosclerotic plaque size. We found that anti-FAP BiTE-treated animals showed a significant increase in the ACTA2^+^ in the fibrous cap within aortic root plaques compared to controls (**Fig. 5h, j)**. These findings suggest that depleting *Fap*^+^ cells with a targeted BiTE may offer a promising therapeutic strategy for mitigating atherosclerotic plaque burden while stabilizing lesions by directly removing modulated SMCs.

**Figure 5.**
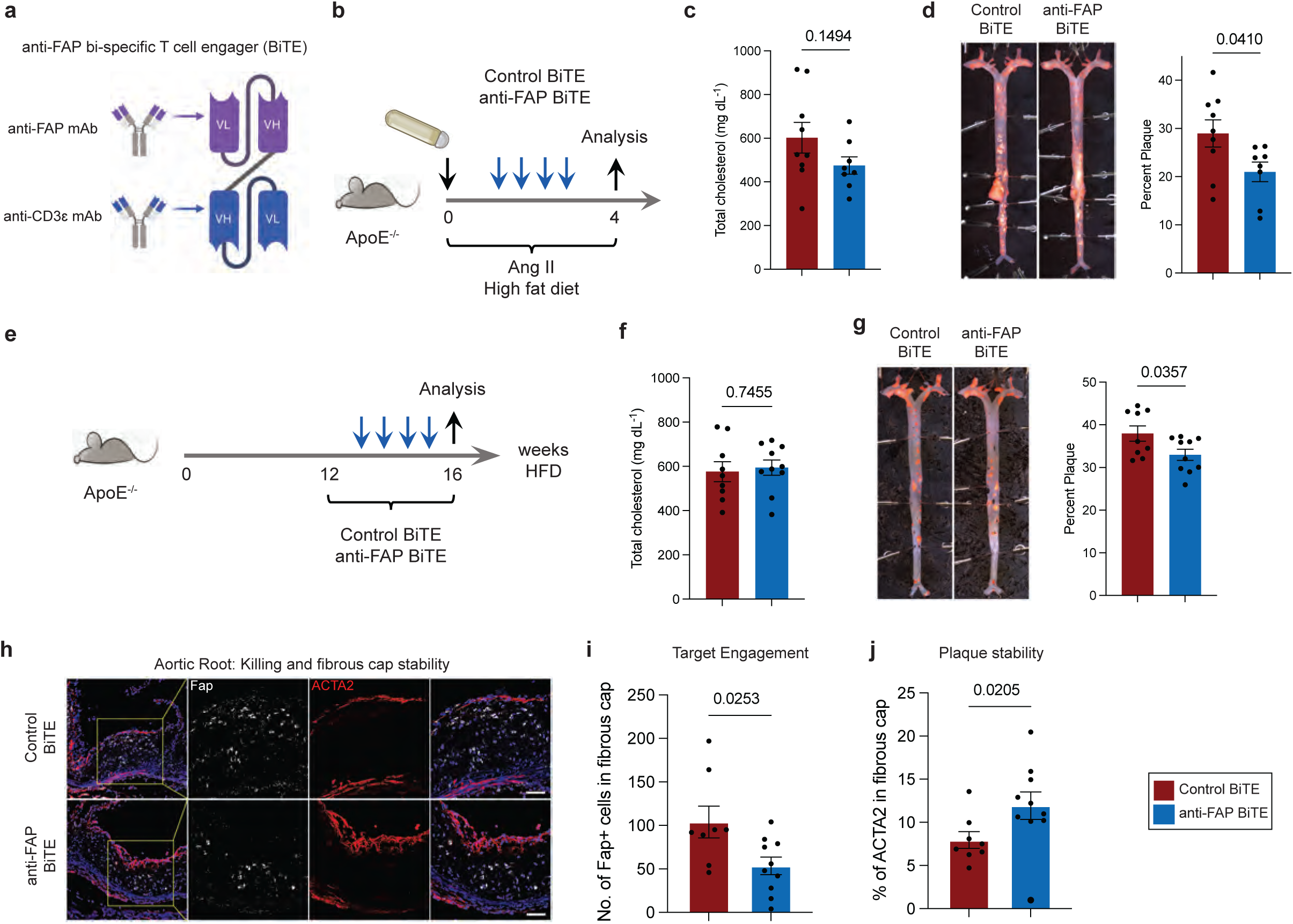
FAP-Targeted Therapy in Murine Atherosclerosis. (a) Schematic of the anti-FAP BiTE construct (Methods). (b) Study design for acute atherosclerosis murine model in *Apoe*^-/-^ mice with HFD feeding/Ang II infusion and control or anti-FAP BiTE treatment for 4 weeks. (c) Plasma cholesterol in control and anti-FAP BiTE treated mice after 4 weeks of HFD feeding; n = 8-9/group (d) En face ORO staining of whole aorta plaque (left) and quantification of plaque lesion in the aortic archfrom control and anti-FAP BiTE treated mice (right); n = 8-9/group. (e) Study design for chronic atherosclerosis murine model in *Apoe*^-/-^ mice with HFD feeding for 16 weeks and control or anti-FAP BiTE intervention for 4 weeks. (f) Plasma cholesterol in control and anti-FAP BiTE treated mice after 16 weeks of HFD feeding; n = 9-10/group (g) En face ORO staining of whole aorta plaque (left) and quantification of plaque lesion in the aortic arch from control and anti-FAP BiTE treated mice (right); n =9-10/group. (h) RNA in situ hybridization for *Fap* and ACTA2 immunofluorescence in aortic root sections from control and anti-FAP BiTE treated animals; Outlined areas indicate the region magnified in the next panels. (i) Number of Fap+ cells in the fibrous cap relative to plaque size. (j) Percent of ACTA2+ area in the fibrous cap; n = 8-10/group. Scale bars, 50 µm. All data were analyzed with unpaired nonparametric Mann-Whitney test. All histological and plaque quantification for in vivo studies was done blinded.

## Discussion

We provide a comprehensive single-cell and spatially resolved atlas of human coronary artery disease (CAD) to dissect the plaque microenvironment and uncover FAP as a key marker of modulated SMCs within the neo-intima. This work builds upon prior studies that characterize the phenotypic landscape and shed light on the epigenetic diversity of SMC subsets within mouse and human atherosclerotic plaques^1,2,5,56–59^ ^14,15,37,39,59–61^. Using single-cell multi-omics, spatial transcriptomics, and in vivo lineage tracing, we establish that FAP⁺ cells originate from medial vascular SMCs and reside within niches that contain lipid associated macrophages. FAP^+^ cell states are highly enriched with genetic risk variants. Furthermore, we demonstrate the translational potential of targeting FAP⁺ modulated SMCs using BiTE technology, which reduced plaque burden in murine models of atherosclerosis. These findings provide new insights into the cellular and molecular mechanisms driving CAD pathogenesis, introduce a novel lipid-independent target, and support the concept that manipulation of modulated SMC states with biologics may serve as an effective therapeutic approach.

High resolution spatial transcriptomics analyses provide unprecedented opportunities to precisely delineate the cellular architecture of CAD. We reveal distinct spatial niches occupied by heterogenous subsets of SMC, immune cell, and fibroblast subset. For example, we identify a cellular niche comprised of FAP⁺ FCMs and lipid-associated macrophages (Mac4) within the media/intima transition that emerges in CAD, highlighting potential interplay between these cell types in neo-intima formation. The enrichment of inflammatory and ECM remodeling pathways within the neo-intima further supports the hypothesis that inflammation and maladaptive SMC transitions contribute to plaque instability. Future studies investigating cell-cell communication events and effector molecules produced within these niches may provide valuable insights in CAD pathogenesis.

BiTE technology has demonstrated remarkable success in oncology^28,29^, particularly in hematologic malignancies. In contrast to CAR-T cell therapies, which require ex vivo manipulation of patient-derived cell products, BiTEs offer an off-the-shelf approach that harnesses endogenous T cells for precise and tunable cell targeting^62–64^. Here, we provide the first evidence that an anti-FAP BiTE can eliminates FAP⁺ cells in an in vivo atherosclerosis model while improving plaque stability, leading to significant plaque reduction. This targeted immunotherapeutic strategy offers a lipid-independent approach to selectively reshape plaque cell composition by directly targeting pro-atherogenic cell populations within the coronary microenvironment. Our findings establish proof-of-concept for immune-mediated clearance of pathogenic SMCs and position BiTEs as a next-generation immunotherapy for CAD, with potential applications extending to other fibroinflammatory cardiovascular diseases.

We reveal the clinical relevance of FAP⁺ modulated SMCs through non-invasive molecular imaging (PET/CT) using an FAPi tracer in patients with CAD. We observed focal FAPi tracer uptake in coronary plaques, suggesting that FAP-targeted imaging may serve as a biomarker for high-risk lesions. It is possible that differential FAPi uptake patterns across anatomical locations and patient groups may provide insights into stable vs. unstable plaques, identify patients at risk for adverse cardiovascular events, serve as a tool to monitor therapeutic responses.

This study is not without limitations. Mechanisms by which FAP+ modulated SMCs are specified remain incompletely defined and likely involve interactions with macrophages. While we demonstrate the efficacy of FAP-targeted BiTE therapy in an experimental atherosclerosis model, its long-term impact on vascular remodeling, immune homeostasis, and systemic inflammation remains to be elucidated. Given the dynamic, heterogeneous nature of atherosclerotic plaques, it is essential to assess whether sustained BiTE activity influences plaque stability, endothelial function, and adaptive immunity over extended treatment. Although FAP expression is enriched in modulated SMCs within plaques, its role in co-existing fibrotic conditions, including cardiac, renal, and hepatic fibrosis warrants further assessment. A comprehensive evaluation of BiTE specificity and toxicity in relevant preclinical models, including humanized systems, will be critical for ensuring safety and efficacy. Furthermore, understanding how BiTEs reshape the atherosclerotic immune landscape will be essential to optimize and effectively translate this approach.

Collectively, our findings provide an integrated single-cell and spatially resolved map of human CAD, establish FAP as a key marker of pathogenic SMC states, and introduce BiTE immunotherapy as a novel strategy for targeting atherosclerotic plaques. By leveraging immune-based therapies to selectively eliminate disease driving cell populations, our study paves the way for precision medicine approaches in cardiovascular disease and highlights the potential for cell-state-specific interventions.

## Acknowledgments

JMA was supported by the American Heart Association Predoctoral Fellowship (826325) and is currently supported by the Washington University School of Medicine Medical Scientist Training Program and Leducq Foundation Network Seed Grant (#20CVD02). KL is supported by the Washington University in St. Louis Rheumatic Diseases Research Resource-Based Center grant (NIH P30AR073752), the National Institutes of Health [R01 HL138466, R01 HL139714, R01 HL151078, R01 HL161185, R35 HL161185], Leducq Foundation Network (#20CVD02), Burroughs Welcome Fund (1014782), and Children’s Discovery Institute of Washington University and St. Louis Children’s Hospital (CH-II-2015-462, CH-II-2017-628, PM-LI-2019-829), Foundation of Barnes-Jewish Hospital (8038-88), and generous gifts from Washington University School of Medicine. YL is supported by the National Institute of Health [R01 HL150891, R35 HL145212, P41 EB025815]. RSYF is funded by Individual Research Grants from the National Medical Research Council (NMRC) of Singapore (MOH-001480-00) and MOE Academic Research Fund (AcRF) Tier 3 (MOE-000333-00). Hannover: Supported by the Deutsche Forschungsgemeinschaft (DFG, Clinical Research Unit KFO 311, FMB and Clinician Scientist Program PRACTIS, JD), the Leducq Foundation (Transatlantic Network Immunofib, FMB, JD), and “REBIRTH – Research Center for Translational Regenerative Medicine” (State of Lower Saxony; FMB). The study was partially supported from an Amgen sponsored research agreement. Study design schematics were created in BioRender.com. We thank the Genome Technology Access Center at the McDonnell Genome Institute at Washington University School of Medicine for help with genomic analysis. The Center is partially supported by NCI Cancer Center Support Grant #P30 CA91842 to the Siteman Cancer Center. This publication is solely the responsibility of the authors and does not necessarily represent the official view of NCRR or NIH.

## Author Contributions

JA, BA, and KL conceived the study. JA drafted the manuscript with assistance from all authors. JA collected coronary arteries. JA and AB isolated cells from human coronary arteries. JA performed 10x CITE-seq cDNA construction for library preparations and sequencing. TY made all libraries for sequencing and generated Visium FFPE data. JA, SH, SM, CJML, YLP, PC, XL, XZ, and performed all computational analysis. CC extracted clinical data. JA, IJ, and SY performed all in vivo experiments. JD and FB generated and processed all human PET/CT imaging data. WLL and JC optimized and developed the in vitro foam cell model. GSH, AK, VC, DP, JB, YL, and ICR provided valuable guidance in in vivo model optimization. MT, SY, and KC assisted with anti-FAP BiTE generation and validation. JA, CK, NOS, RK, RSYK, TQ, SJ, CML, BA, and KL assisted with data interpretation of multi-omic data. All authors contributed to the experimental design, data analysis and interpretation as well as manuscript production. KL is responsible for all aspects of this manuscript including experimental design, data analysis, and manuscript production. All authors approved the final version of the manuscript.

## Materials and Methods

### Ethical Approval for Human Specimens

The study is compliant with all relevant ethical regulations and has been approved by the Washington University School of Medicine Institutional Review Board (IRB #201104172). Informed consent was obtained from each patient prior to tissue collection by Washington University School of Medicine and no compensation was provided in exchange for subject participation in the study. All demographic and clinical data has been de-identified and provided in **Supplementary Table 1.** Patients included in this study span diverse race, age, and sex to provide an inclusive trans-ethic study population.

### Inclusion Criteria

Prior to tissue collection, specific inclusion criteria were employed to ensure well controlled study groups. Any patients with HIV or hepatitis and known genetic cardiomyopathies were excluded from this study. The left anterior descending coronary artery was isolated, and flash frozen from donor hearts: patients with stable ejection fractions, no known history of cardiac disease and experienced a non-cardiac cause of death/transplant and from patients with chronic heart failure. For all samples the proximal left anterior descending coronary artery was used.

### Human single cell isolation for CITE-seq

Fresh human coronary arteries from ex-planted hearts at the time of transplantation or donors declared DCD at Washington University School of Medicine and Mid America Transplant Service were harvested, perfused with cardioplegia, and transported on ice. Starting at the coronary sinus, the left main, left anterior descending, and left circumflex arteries were dissected removing any epicardial fat while preserving the adventitia. Sections were partitioned for histopathology, spatial transcriptomics, single cell isolation, and flash frozen. Cells were isolated as before^19^. Briefly: Coronaries were minced using a razor blade on ice and transferred to a 15 mL conical tube containing 3 mL DMEM with 170 µL Collagenase IV (250 U mL^-1^ final concentration), 35 µL DNAse1 (60 U mL^-1^), and 75 µL Hyaluronidase (60 U mL^-1^, all purchased from Sigma Aldrich) and incubated at 37 °C for 45 min with agitation. After 45 min, the digestion reaction was quenched with 6 mL of HBB buffer (2% FBS and 0.2% BSA in HBSS), filtered through 40µm filters into a 50 mL conical tube and transferred back into a 15 mL conical tube to obtain tighter pellets. Samples were then spun down at 4°C, 1,200 rpm for 5 min and the supernatant was discarded. Pellet resuspended in 1 mL ACK Lysis buffer (#A10492-01, Gibco) and incubated at room temperature for 5 min followed by the addition of 9 mL DMEM and centrifugation at 4°C, 1,200 rpm for 5 min. Supernatant was discarded and the pellet was resuspended in 2 mL FACS buffer (2% FBS and 2 mM EDTA in calcium/magnesium free PBS) and centrifugation was repeated in above conditions and supernatant aspirated. The TotalSeq A 277 panel (BioLegend) antibody cocktail was resuspended in 100 uL of FACS buffer and 1 µL each of custom oligo-labeled FAP (Amgen) and LRRC15 (Amgen) were added. The combined 102 µL were used to resuspend the pellet with the addition of 1 µL of DRAQ5 (#564907, Thermo Fisher Scientific) and incubated on ice for 30 min. Solution was washed with FACS buffer three times following same centrifugation as above and then resuspended in 500 µL of FACS buffer and 1 µL DAPI (#564907, BD Biosciences,) and filtered into filter-top FACS tubes. First singlets were gated and subsequent DRAQ5+/DAPI-events were collected in 300 µL cell resuspension buffer (0.04% BSA in PBS) – collected cells were centrifuged as above and resuspended in collection buffer to a target concentration of 1,000 cells µL^-1^. Cells were counted on a hemocytometer before proceeding with the 10x protocol.

### CITE-seq Library Preparation

Collected cells were processed using the single Cell 3’ Kit v 3.1 (#1000268, 10x Genomics). 10,000 cells were loaded onto ChipG (#1000121) for GEM generation. Reverse transcription, barcoding, and complementary DNA amplification of the RNA and ADT tags were performed as recommended in the 3’ v3.1 chromium protocol. Custom conjugated FAP antibodies were generated using TotalSeqA barcodes by Biolegend. sequenced on a NovaSeq6000.

CITEseq libraries were generated using TotalSeqA reagents and Chromium Next GEM Single Cell 3ʹ Kit v3.1 reagents (#1000268, 10x Genomics) following the 10x Genomics protocol (Document CG000185) modified to capture the TotalSeqA barcodes. Briefly, during cDNA amplification, 1 µl of 0.2 µM ADT Additive Primer (CCTTGGCACCCGAGAATT*C*C) was added to the reaction. 5 µl of purified ADT fraction was indexed with SI-PCR primer (AATGATACGGCGACCACCGAGATCTACACTCTTTCCCTACACGACGC*T*C) and RPI Primer (CAAGCAGAAGACGGCATACGAGAT[i7index]GTGACTGGAGTTCCTTGGCACCCGAGAATT C*C*A) at a final concentration of 2.5 µM each using 2x Kapa Hifi Master Mix. Libraries were sequenced on a NovaSeq 6000.

### CITE-seq alignment, quality control, and cell type annotation

Raw fastq files were aligned to the human GRCh38 reference genome using CellRanger (10x Genomics, v6.1) with the antibody capture tag for the TotalSeqA 277 + two custom oligo-tagged antibodies. First, protein reads were normalized and de-noised for each sample separately using the dsb package^65^ in R v4 with the isotype controls tag to remove noisy cell-to-cell variation. Subsequent quality control, normalization, dimensional reduction, and clustering was performed in Seurat v4.0. Following normalization, quality control was performed and cells passing the following criteria were kept for downstream processing: 500 < nFeature_RNA < 600 and 1,000 < nCount_RNA < 25,000 and percentage mitochondrial reads < 15%. Raw RNA counts were normalized and scaled using SCTransform^66^ regressing out percent mitochondrial reads and nCount_RNA. Principal component analysis was performed on normalized RNA counts and to determine the number of PCs to use for further processing the following criteria was applied: PCs exhibit cumulative percent > 90% and the percent variation associated with the PCs as < 5%. Weighted nearest neighbor clustering^67^ (WNN) was performed with the significant RNA PCs and dsb normalized proteins (minus isotype controls) directly without PCA as previously outlined with the FindMultiModalNeighbors function in Seurat. Subsequently, a uniform manifold approximation (UMAP) embedding was constructed and FindClusters was used to un-biasedly cluster cells using the SLM modularity optimization algorithm. Clustering was performed for a suite of different resolutions (0.1-0.8 at 0.1 intervals) and differential gene expression using the FindAllMarkers function and a Wilcoxon Rank Sum test with a logFC cut-off of 0.58 and a min.pct cut-off of 0.1. Clusters were annotated using canonical gene and protein markers and subsequent violin plots (RNA) and heatmaps (protein) were created to assess clean separation of clusters into distinct cell types. Previously identified canonical marker genes were also plotted on the UMAP object to further validate cluster annotations.

### Spatial transcriptomics data generation

Visium FFPE: Fresh coronary tissue was fixed in 4% PFA overnight, washed with PBS, and embedded in FFPE blocks. FFPE blocks were sectioned onto Visium slides and underwent H&E staining before imaging. Libraries were generated using the Visium FFPE v1 kit (#1000188; #1000362; #1000364) and sequenced on a NextSeq2000 according to manufacturer’s recommendations. Libraries were sequenced on a NovaSeq 6000. Xenium in situ: A custom gene panel was designed (**Supplementary Table 4**) based on the CITE-seq reference map and combined with the Xenium Human Multi-Tissue and Cancer Panel (#1000626, 10x Genomics). FFPE blocks and sectioned and the 10x Genomics Xenium In Situ Gene Expression with Cell Segmentation Staining User Guide protocol (10x Genomics, Protocol number CG000749) used for sample (#1000661; #1000460; #1000487. 10x Genomics,).

### Mouse to human integration

Published murine atherosclerosis data was internalized and re-processed using the methodology described above. We used the convert_mouse_to_human_symbols function within the nichenetr^68^ library to convert mouse genes to the corresspoding human gene name. FindTransferAnchors and MapQuery functions were used to reference map the mouse data onto the human CITE-seq map using a superviced PCA as the reference reduction.

### Spatial transcriptomics computational analysis: Visium FFPE

Visium FFPE data was aligned using space ranger and used for downstream analysis and voxels containing no transcripts were included in the downstream analysis. Data was normalized using SCTransform with nCount_Spatial regressed out. We used the FindVariableFeatures to identify 3000 highly variable genes and the runNMF function was used from the GeneNMF package^51^ with ndim=30 to decompose the data into interpretable gene programs. To account for batch effects across samples, we applied multiNMF (k=4:9 and nfeatures = 3000) to combine gene programs into consensus MPs that were conserved across samples. We then represented each spatial voxel by its MP score and used these components to generate a UMAP visualization, with nearest-neighbor clustering, to identify conserved spatial niches across samples. Differential gene expression analysis revealed key marker genes associated with these spatially conserved niches. 10X visium spatial colocalization analysis: tangram^36^ was used to deconvolute cell-types and cell-states for the 16 samples. The probability prediction of every spot being a specific cell type is obtained as an output from tangram analysis. This spot-level output from tangram was used as the input to predict spatial localization. Specifically, the Misty function, implemented in liana (1.4.0) was used for colocalization per-slide using a random forest model (default parameters).

### Spatial transcriptomics computational analysis: Xenium

Raw sequencing data is aligned and counts decoded into spatial transcripts. The cellular stains (nuclear, cell interior, cell boundary) and a neural network algorithm is used for cell segmentation and transcripts are assigned to the segmented cells. This yields a cell by gene matrix with corresponding spatial coordinates for each cell. This matrix is used for downstream analysis including nearest neighbor analysis, clustering, and UMAP embedding construction. We then use differential expression analysis to annotate clusters into cell types and perform reference mapping to a single cell atlas to validate annotations. High-resolution spatial colocalization analysis: The Xenium data tangram output was obtained for sixteen distinct slides, adding predictions for every spot. Using the random forest model, the liana^69^ (1.4.0) wrapper’s misty() function conducts misty analysis for every slide. The co-localization of various cell types was ascertained using the importance score values between target and predictor. Lumina to Periphery: Lumina to periphery cell-type distribution was calculated in all arteries^70^. Briefly, for each Xenium slide, first the centre of the artery was estimated using mean spatial coordinates. Then the normalised distance between the center to each spot in the slide was calculated. The spots with minimal and maximal distance were marked in the entire slide by varying the angles from the center(-180 to +180) to cover 360 degree round, in order to define the lumina and periphery. Each spot was defined as a value based on the closest and farthest points to define the lumina to periphery gradient. Niches discovery in high resolution spatial data: To determine groups of spots sharing similar compositions of cell types, the proportion of cell type distribution from tangram output was converted to isometric log ratios. This clustered the spots into different niches sharing similar cellular compositions. Louvain clustering of spots was calculated by running the nearest neighbour graph on different k numbers (10, 20, 50). The clustering resolution was determined using the value that maximizes the mean silhouette score of each cluster. The over-representation of cell types in each niche was determined by comparing the distribution of cell types in the niches vs. the rest using the Wilcoxon test (pval<0.05).

### FAPi-PET/CT acquisition

FAPi-targeted PET/CT (positron emission computed tomography / computed tomography) was conducted in 10 patients with acute myocardial infarction (AMI) and 5 patients with chronic ischemic cardiomyopathy (ICM) using the specific ligand 68Ga-FAPI-46, which was synthesized in house according to good manufacturing practice, as previously described^41^, and used clinically according to §13.2b of the German Pharmaceuticals Act, for determination of myocardial profibrotic activity. In patients with AMI, PET was acquired 7.9±1.6 days after myocardial injury. Dual gated (respiratory and ECG-gating for improved cardiac imaging) PET images were acquired for 20min using a Biograph mCT 128 system (Siemens, Knoxville, TN, USA), beginning 60min after intravenous injection of 130±41 MBq of 68Ga-FAPI-46. Images were iteratively reconstructed, using time-of-flight and point-spread function information (True X, Siemens). Image interpretation and acquisition of standardized uptake values (SUV) was conducted using commercial software (syngo.via; V50B, Siemens Healthcare).

### FAPI-PET/CT analysis

PET/CTs were systematically analyzed to assess focal coronary arterial uptake of the specific FAPI-tracer Ga68-FAPI-46. Visual assessment determined the presence or absence of uptake in regions corresponding to native or treated coronary lesions (experienced reader JD). These lesions were defined based on the presence of coronary stents or calcifications detected via CT imaging. Following this visual evaluation, coronary lesions were confirmed using cardiac catheterization reports to ensure the accuracy of the identified lesions.

### BiTE^TM^ molecules

For this study, BiTE HLE molecules were constructed with a mouse immunoglobulin G1 (IgG1) Fc domain engineered with scaffold lacking effector function (SEFL1) mutations^71^. These mutations greatly reduce Fcg receptor binding and abrogate antibody effector functions such as antibody dependent cellular cytotoxicity and antibody-dependent cellular phagocytosis. The following BiTE molecules were used in this study: (A) 64281 muFAPa BiTE containing a mouse anti-mouse FAPa scFv derived from the 73.3_MABC1145 antibody; and (B) 64280 control BiTE containing an anti-idiotype scFv that does not recognize a known target derived from the Amgen mouse mAb E8.1. The above BiTE molecules also contain the KE3 BiTE^72^ consisting of a mouse anti-mouse CD3e KE3 scFv derived from the KT3 mAb sequence^73^. BiTE HLE molecules were produced using standard protocols described previously^74^. Briefly, BiTE HLE molecules were expressed from CHO stable cell lines using serum-free media and purified from cell culture supernatant using protein A capture (Mabselect Sure, Cytiva) followed by preparative size-exclusion chromatography (Superdex 200, Cytiva). Final product quality was confirmed by mass spectrometry (ThermoFisher Vanquish LC / Q Exactive Plus Orbitrap), analytical size exclusion chromatography (HPLC-SEC, Agilent 1100 Series HPLC) and endotoxin testing (EndoSafe®-MCS).

### Diet and assessment of atherosclerosis

All experimental mice were fed a high fat diet containing 21% fat by weight (42% kcal from fat) and 0.2% cholesterol (#TD.88137, Envigo Teklad) for 4, 16 (For *Apoe^-/-^*) and 24 weeks (For *Myh11*^creERT2^ Rosa26^tdTomato^) starting at 8 or 9 weeks of age. To activate Cre-recombinase, *Myh11*^creERT2^ Rosa26^tdTomato^ mice were gavaged with 2 mg of tamoxifen (#T5648, Sigma-Aldrich) in 0.1 ml of corn oil (#C8267, Sigma Aldrich) for 5 consecutive days starting at 6 weeks. After 2 weeks of recovery, mice were intravenously injected with 5 × 10^11^ vector genome copies of AAV8-D377Y-mPCSK9 (Vector Biolabs, United States) at 9 weeks of age and immediately placed on an HFD. After HFD feeding, blood was collected from the retro-orbital plexus. Mice were euthanized by carbon dioxide inhalation. Plasma samples were prepared from the collected blood by centrifugation at 13,000 rpm for 10 min at 4°C. Total cholesterol in plasma were determined using the appropriate kit (#C7510-500, Pointe Scientific, Inc). Hearts and whole aortas (from the aortic arch to the iliac artery) were harvested after perfusion with PBS. For en face analysis, isolated aortas were cleaned by removing perivascular fat tissues, opened longitudinally, and pinned onto black wax plates. After fixation with 4% paraformaldehyde overnight at 4°C, aortas were washed with PBS for 1 hr, and stained with 0.5% Oil Red O in propylene glycol (#O1516, Sigma-Aldrich) for 3 hr at room temperature. After staining, aortas were de-stained with 85% propylene glycol in distilled water for 5 min to reduce background staining and washed with distilled water for 15 min. For analysis of plaque in the aortic root, hearts were fixed overnight with 4% paraformaldehyde at 4°C, washed with PBS for 1 hr, and embedded into optimal cutting temperature (OCT) compound (#4583, Sakura Finetek). 5-µm-thick cryosections were used for immunofluorescent studies, and stained area were digitized and calculated using AxioZen or AxioVison (Carl Zeiss).

### In situ hybridization (ISH)

To detect RNA transcripts for Fap in both human and mouse vascular tissues, a commercially available kit (#323100, RNAscope Multiplex Fluorescent Reagent Kit v2, Advanced Cell Diagnostics) was used according to the manufacturer’s instructions. Briefly, 4% PFA-fixed human and mouse frozen sections with 5-µm-thickness were air-dried for 1 hr at room temperature and treated with hydrogen peroxide for 10 min to block endogenous peroxidase activity. After antigen retrieval by boiling in target antigen retrieval solution for 5 min at ∼100°C, slides were treated with protease III for 30 min at 40°C. Target probes (#423881, mouse Fap) were hybridized for 2 hr at 40°C, followed by a series of signal amplification and washing steps. Hybridization signals were detected by TSA Plus Cyanine 5, and co-stained with anti-Col1A1XP-Alexa488 (#28368S, Cell Signaling).

### Sample preparation for single-cell RNA sequencing of foam cells

THP-1 cells (#TIB-202, ATCC;) were cultivated in RPMI-1640 medium (#10-041-CV, Corning;) with 10% FBS. They were then treated with PMA (#5.00582.0001, Millipore Sigma) to reach 15 nM concentration and plated in a 12-well plate (#150628, Thermos Fisher Scientific) at a density of 5 x 10^5^ cells per well. The plate was incubated for a duration of 3 days to promote differentiation. On the fourth day, the cells were gently washed with 1 mL of fresh medium to remove any residual PMA and other soluble factors. Following this, oxLDL (#L34357, Invitrogen) was introduced to each well to reach a final concentration of 50 µg mL^-1^, ensuring thorough mixing for uniform exposure. The cells were incubated for an additional four days to allow for foam cell formation. Upon completion of the incubation period, the medium was carefully aspirated from each well. The cells were detached by adding 200 µL of trypsin-EDTA (#25-053-CI, Corning) per well and incubating at 37°C for 5 min to facilitate cell detachment. To neutralize the trypsin activity, 1 mL of medium was added per well and the mixture was gently pipetted to ensure complete suspension of the cells. Finally, the cell solution was filtered through a 100 µm nylon cell strainer (#352360, Falcon) to remove any cell clumps, thereby obtaining a uniform single-cell suspension ready for further analysis.

### RNA extraction and reverse transcription quantitative PCR

Total RNA was isolated from the cells by using TRIzol Reagent and the RNeasy Mini Kit (#74104, QIAGEN;). To lyse the cells, 1mL of TRIzol Reagent was added and homogenized by pipetting the lysate several times. The mixture was incubated for 5 min following by adding 0.2 mL of chloroform. The mixture was shaken thoroughly and incubated for 3 min. After centrifugation at 12,000g for 15 min at 4°C, the top RNA-containing aqueous phase was collected and extracted via QIAcube machine with RNeasy mini standard protocol (#79254, QIAGEN) and additional on-column DNase digestion step. The qPCR reactions were performed using TaqMan RNA-to-CT 1-Step Kit in QuantStudio 7 Flex Real-Time PCR Systems. Primers were purchased from Thermo Fisher. The fold-change in relative gene expression was calculated using the 2^−ΔΔCT^ method. The ribosomal RNA18S5 was used as internal control.

### OxLDL uptake image analysis

THP-1 cells were exposed to 50 nM PMA at 37°C for three days, followed by an 18-hr incubation with a corresponding concentration of Dil-oxLDL (#L34358, Invitrogen;). The medium was then removed, and the cells were fixed in 4% PFA at room temperature for 15 min. After fixation, the cells were treated with 1 µg ml^-1^ Hoechst 33342 (#BDB561908, BD Biosciences) and 1 µg ml^-1^ WGA488 (#W11261, Thermos Fisher Scientific) in PBS for 1 hr at room temperature. Subsequently, the PBS was refreshed, and images were captured using a fluorescence microscope (Opera Phenix High-Content Screening System, Revvity). The population of cells with oxLDL is quantified by normalizing the number of Dil-oxLDL cells to the number of nuclei.

### Oil red O and hematoxylin staining

The Oil Red O staining kit (#MAK194, Sigma Aldrich) was used. Cells were fixed in 4% PFA for 30 min and then rinsed with water twice. Subsequently, the cells were treated with 60% isopropanol for 5 min before being covered evenly with Oil Red O Working Solution for a duration of 15 min. The staining solution was then discarded, followed by washes with water five times. Cells were incubated with hematoxylin for 1 min, after which they were washed again with water five times. Finally, the cells were immersed in water and examined under a microscope (EVOS imaging system).

### 10x scRNA-seq data analysis of in vitro foam cells

Initial processing of the 10x scRNA-seq data was done with the Cell Ranger Pipeline (v.7.1.0) by first running cellranger mkfastq to demultiplex the bcl files and then running cellranger count with 10x Genomics’ pre-built Cell Ranger reference GRCh38-2020-A_build. After running Cell Ranger, the filtered_feature_bc_matrix was read into R (v.4.2.1) with the Seurat (v.5.1.0) function Read10X^75^. Doublets were detected by scDblFinder (v.1.2.0) for each sample separately and removed from the downstream analyses^76^. Cells from all samples were merged into a single Seurat object and analyzed separately for each dataset. Cell barcodes were filtered based on the number of genes per cell (between 1,000 and 8,000), unique molecular identifiers (UMIs) per cell (between 2,000 and 50,000), and percentage of mitochondrial reads per cell (less than 10%). The filtered cell number used for subsequent analysis was 9,397 for the THP-1 culture, 9,016 for the THP-1 + 15 nM PMA culture, 13,781 for the THP-1 + 15 nM PMA + 50 µg ml^-1^ oxLDL culture, and 9,298 for the THP-1 + 15 nM PMA + 50 µg ml^-1^ Dil-oxLDL culture, respectively. Read counts were normalized using Seurat’s LogNormalize method. Cell-cycle scoring was calculated with the CellCycleScoring algorithm from Seurat using cell-cycle-related genes and the difference between the G2M and S phase scores were regressed out. The 2,000 most variable genes were identified for subsequent analysis. Principal component analysis (PCA) was performed using the RunPCA function and the top 30 principal components (PCs) were selected. Batch effect was removed using Harmony (v. 0.1.1)^32^. UMAP was computed using the RunUMAP function and cells were clustered using a k-nearest neighbour graph and the Leiden algorithm with the FindNeighbors and FindClusters functions. A resolution of 0.9 was used for clustering. The FindAllMarkers and FindMarkers functions were performed after normalizing the expression counts with Seurat’s “LogNormalize” method to identify the marker genes for each cluster (min.pct = 0.1, Padj < 0.05, logfc.threshold = 0.25) and intercluster differentially expressed genes (min.pct = 0.25, Padj < 0.01, logfc.threshold = 0.5) via the Wilcoxon rank sum test. Pathway analysis of marker genes and differentially expressed genes was conducted with QIAGEN IPA (QIAGEN Inc., https://digitalinsights.qiagen.com/IPA).

### Integration of human coronary myeloid 10x scRNA-seq datasets

We performed reference mapping to project the THP-1 derived foam cells onto the UMAP structure of the 10x scRNA-seq data of the human coronary myeloid cells using Seurat. Specifically, the FindTransferAnchors function was used to find a set of anchors between the reference and query object with the top 30 PCs and “LogNormalize” normalization. These anchors were then passed into the MapQuery function to project the query data onto the UMAP embedding of the reference.

### Statistics and Reproducibility

No sample size calculations were performed. Sample size was governed by tissue availability and input tissue mass was based on ability to recover sufficient cells. No samples were excluded. In vivo experiments were replicated at-least once for a total of two batches and data was pooled for downstream analysis (all attempts at replication were successful). For human studies all transplants with HF, MI, and non-failing donors were processed randomized across age, sex, and race. For in vivo studies male mice were used and randomized to control and treatment groups. All histological and plaque quantification for in vivo studies was done blinded. All single cell/nuclei analysis including clustering is done using un-biased techniques. Investigators were blinded to the groups when performing initial analysis. Blinding during data collection was not necessary as nuclei isolation protocol required FACS to collect intact cells/nuclei with no exclusion of any cells/nuclei. This sorting approach does not introduce any bias into the sample collection.

## Data Availability

Raw and processed sequencing files can be found on the Gene Expression Omnibus super series ().

## Code Availability

Scripts used for analysis in this manuscript can be found at (https://github.com/jamrute).

**Supplementary Figure 1.** (a) Study design with isolation of single cells from human coronary arteries using FACS for DRAQ5^+^/DAPI^-^ gating. (b) Quality control metrics for RNA and ADT CITE-seq data post-QC. (c) Scrublet score in UMAP plot after doublet removal. (d) Cell type composition across all samples.

**Supplementary Figure 2.** (a) Number of spots under tissue (left) and genes detected (right) across samples from Visium FFPE. (b) Quality control metrics across samples post-QC. (c) H&E staining image of all human coronary arteries used for Visium FFPE processing. (d) Tangran deconvolution scores for major cell types (SMC/Pericyte, modSMC, endothelium, Fibroblast1, myeloid, and T/NK cells in healthy, obstructed, and ruptured plaques.

**Supplementary Figure 3.** (a) Xenium quality control metrics across annotated cell types post-QC. (b) Zoom in within the intima showing 4-color stains and segmented cells colored by imputed annotation in Fig. 1b. For differential gene and protein expression, the Wilcoxon Rank Sum test is used to adjust the p-value. (c) UMAP plot of Xenium segmented cells annotated from CITE-seq reference mapping (Fig. 1b) split by plaque burden. (d) Example of mild, moderate, and severe plaque Xenium sections colored by reference mapped cell type annotations.

**Supplementary Figure 4.** (a) DotPlot of stromal cell state marker genes from Fib. 2a; for differential gene expression, the Wilcoxon Rank Sum test was used to adjust the p-value. (b) Genome-wide Spearman correlations between FAP and all other genes (in Pericytes, SMC1-4, FMC, and CMC) with ranking of top 30 that are statistically significant; multiple testing correction using Benjamini-Hochberg method. (c) *MGP* gene expression in stromal cell UMAP plot. (d) Spearman correlations between FAP and all other proteins (in Pericytes, SMC1-4, FMC, and CMC) in CITE-seq panel with ranking of all statistically significant proteins; multiple testing correction using Benjamini-Hochberg method. (e) ITGA2 protein expression in stromal cell UMAP plot. (f) Gene Ontology biological processes pathway enrichment for genes correlated with FAP protein levels; the adjusted p-value is computed using the Bejamini-Hochberg method for correcting for multiple hypotheses testing.

**Supplementary Figure 5.** (a) Heatmap of genes enriched along the contractile to FMC/CMC Palantir trajectory with key genes highlighted. (b) Reference mapped murine atherosclerosis data onto human CITE-seq UMAP. (c) Heatmap of average cell prediction score of human CITE-seq cell types (y axis) in annotated mouse cells (x axis). (d) Ridge plot for mapping score distribution across all cell types. (e) Endothelium (*Cdh5*) and SMC (*Myh11*) cell labeled cells mapped onto human CAD CITE-seq data colored by cell type. (f) Heatmap of average cell prediction score of human CITE-seq stromal cell states (y axis) in annotated mouse *Myh11* lineage positive cells (x axis). (g) Study schematic of *Myh11* lineage tracing data in murine atherosclerosis from published work. (h) *Fap* expression grouped by time-point post-HFD feeding of tdTomato^+^ lineage cells. (i) RNA in situ hybridization for *Fap* and COL1A1 immunofluorescence in aortic root sections from *Apo*e^-/-^ mice after 16-weeks of HFD feeding; Outlined areas indicate the region magnified in the next panels.

**Supplementary Figure 6.** (a) MP pairwise similarity in terms of Jaccard index between constituent gene programs which make up consensus MPs. (b) DotPlot of differentially expressed genes across spatially conserved Niches; for differential gene expression, the Wilcoxon Rank Sum test is used to adjust the p-value. (c) MP gene set score overlaid on UMAP plot of spatial niches. (d) Heatmap of MP gene set score across tangram deconvolute cell type in each voxel. (e) Spatial niches overlaid on Visium FFPE spatial sections across lesion severities; colored by Niches in Fig. 3(a). (f) PROGENy pathway enrichment analysis for NF-κB in a healthy (left) and diseased coronary artery. (g) FAP module gene set score overlaid on a healthy (left) and diseased coronary artery.

**Supplementary Figure 7.** (a) UMAP plot of Xenium stromal cells colored by reference mapped annotations from CITE-seq stromal map (Fig. 2a). (b) Example Xenium images of stromal cell states annotated in space. (c) *MGP*, *VCAN*, and *LTBP2* gene expression in Xenium sections.

**Supplementary Figure 8.** (a) DotPlot of myeloid cell state marker genes from Fib. 4c; for differential gene expression, the Wilcoxon Rank Sum test is used to adjust the p-value. (b) Surface protein expression for LILRA4, CD276, CCR2, and CD28 in myeloid UMAP plot. (c) Expression of myeloid specific genes linked to CAD GWAS variant across myeloid cell states. (d) *LIPA* gene expression in a Visium FFPE section from healthy (top) and diseased (bottom) coronary artery. (e) FAP module (left) and Mac4 gene set score in a diseased coronary artery with white arrow highlight overlap.

**Supplementary Figure 9.** (a) Oil-red-O staining in THP-1 cells treated with PMA and THP-1 cells treated with PMA and oxLDL. (b) Quantification of the percentage of cells with oxLDL uptake in treatment of different concentrations of oxLDL. (c) Quality control metrics by condition of in vitro LAMs. (d) UMAP plot of in vitro macrophage cell states with oxLDL treatment. (e) DotPlot of IVMac cell state marker genes; for differential gene expression, the Wilcoxon Rank Sum test was used to adjust the p-value. (f) Cell state composition of IVMac cell states across conditions. (g) Density plot of Mac4 gene set score from the human CAD CITE-seq map (Fig. 4c) overlaid on IVMac UMAP.

