## Supplementary figures and images for "Targeting Modulated Vascular Smooth Muscle Cells in Atherosclerosis via FAP-Directed Immunotherapy"

### Supplemental Figure 1

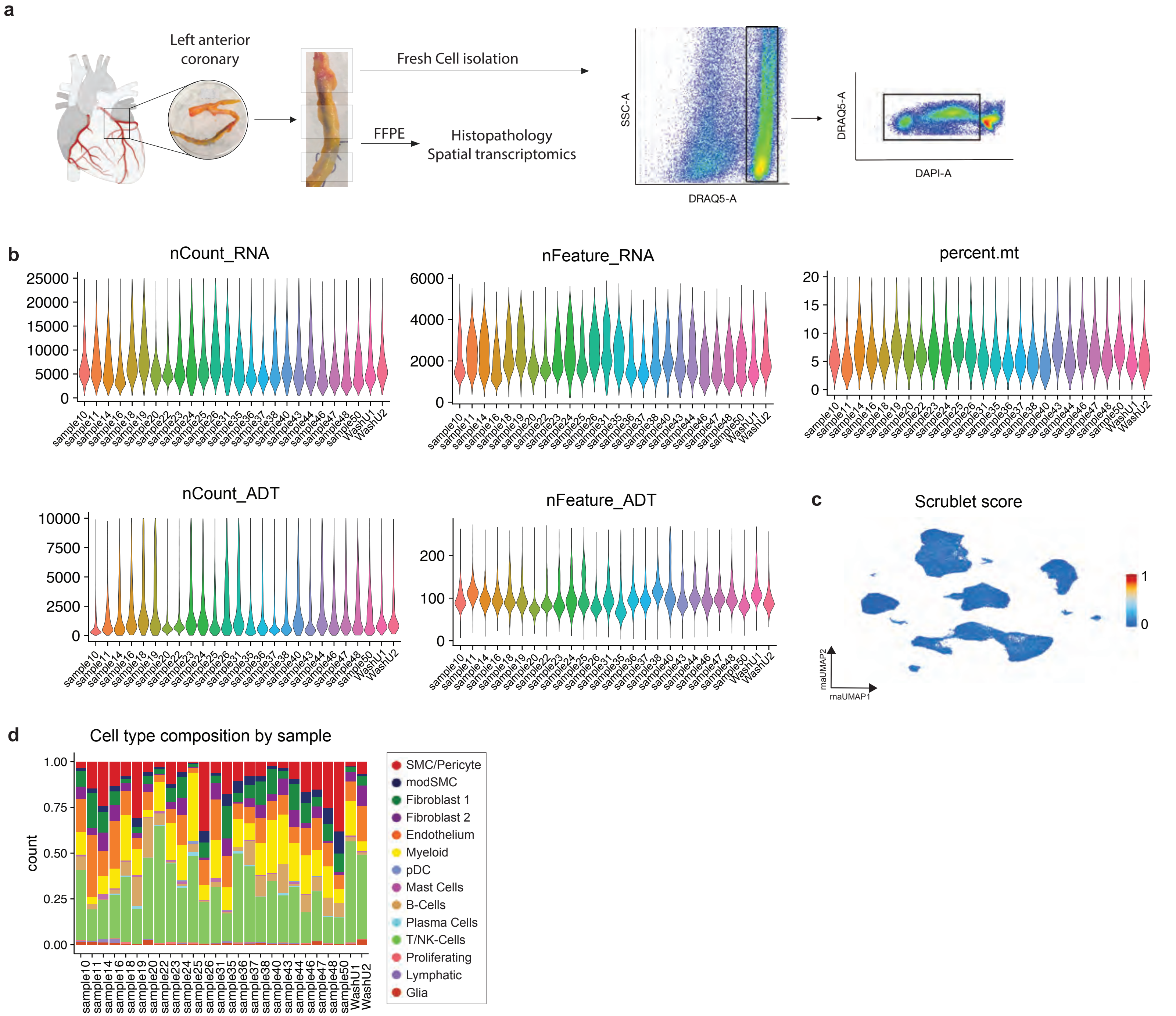

### Supplemental Figure 2

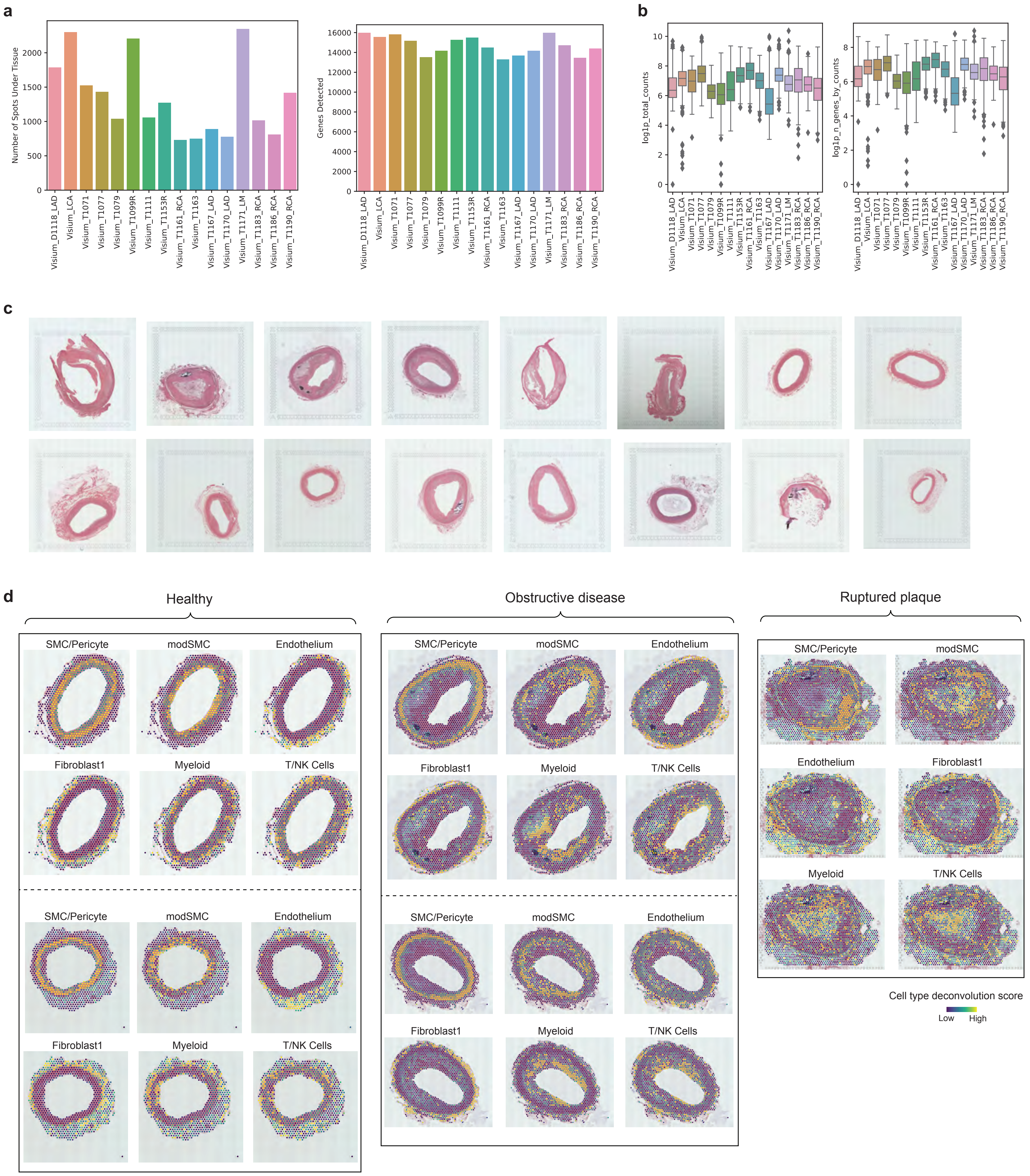

### Supplemental Figure 3

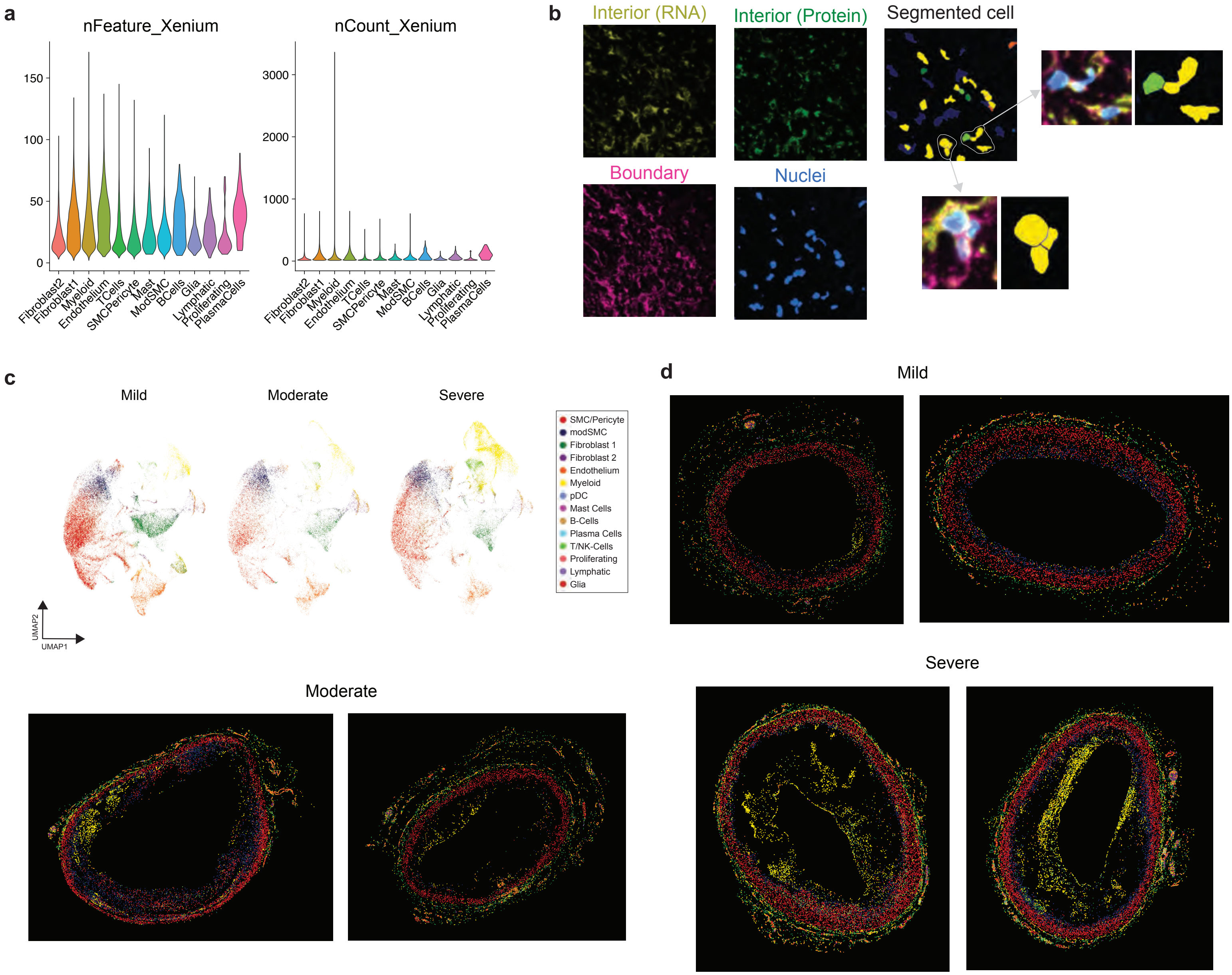

### Supplemental Figure 4

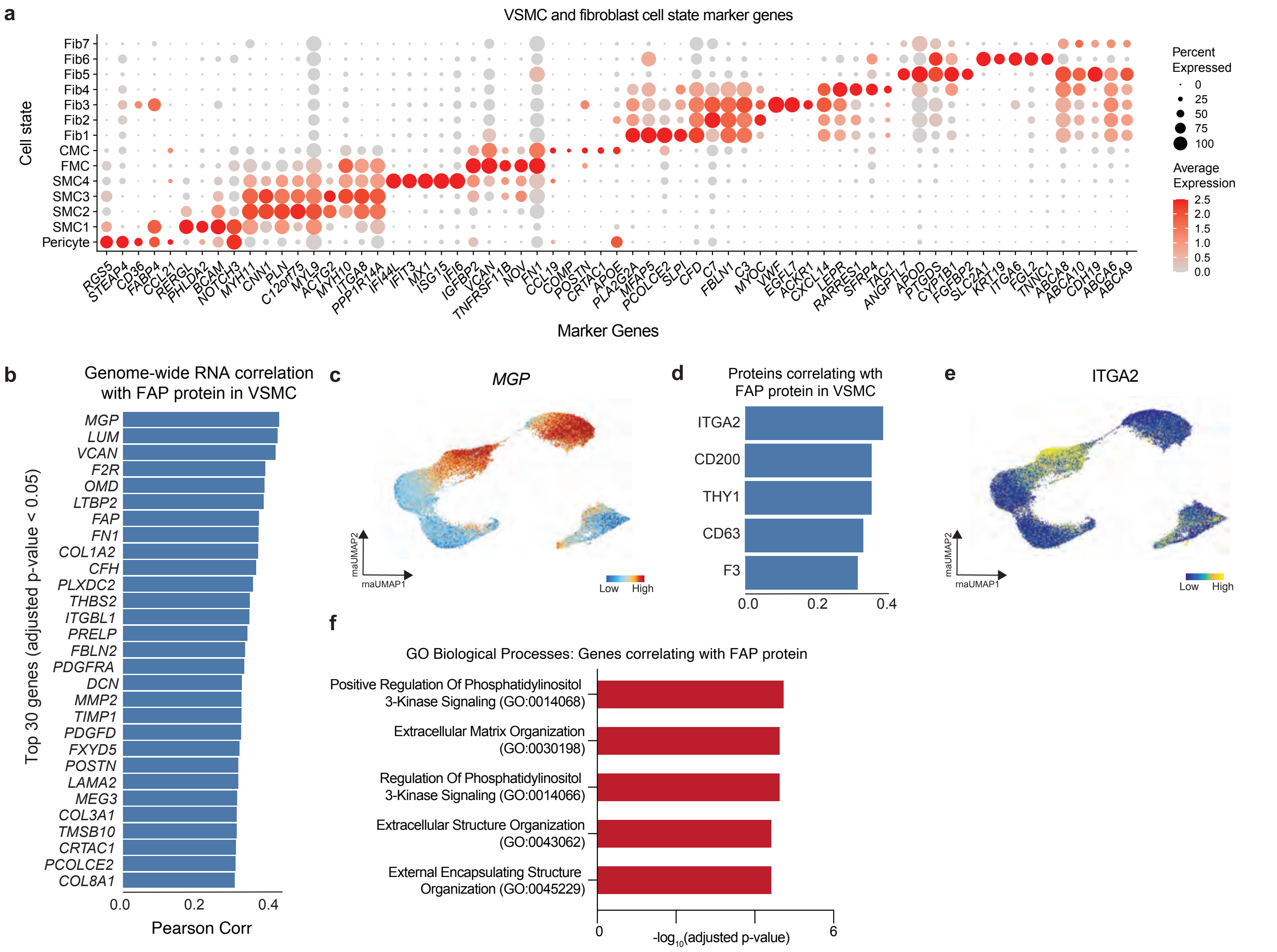

### Supplemental Figure 5

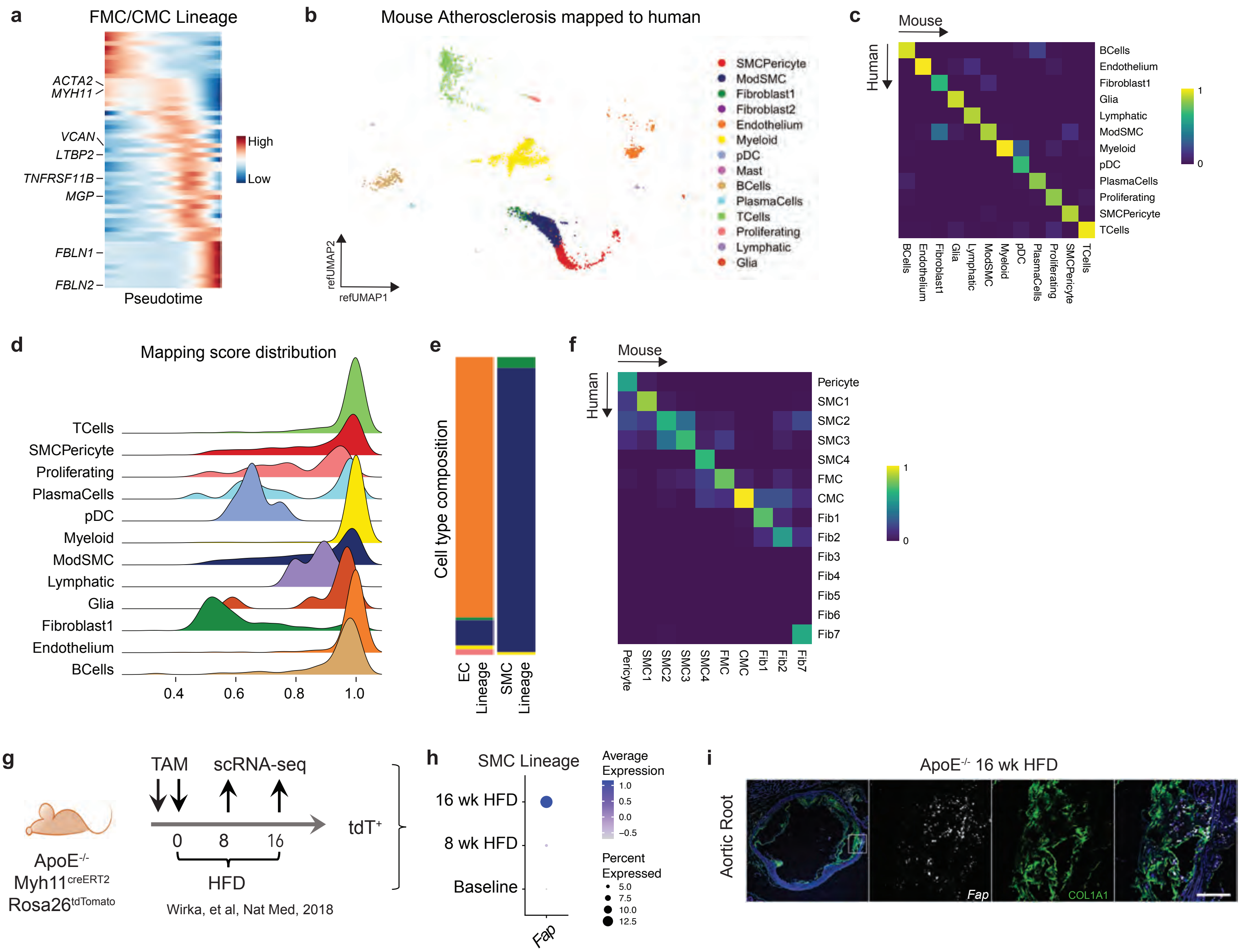

### Supplemental Figure 6

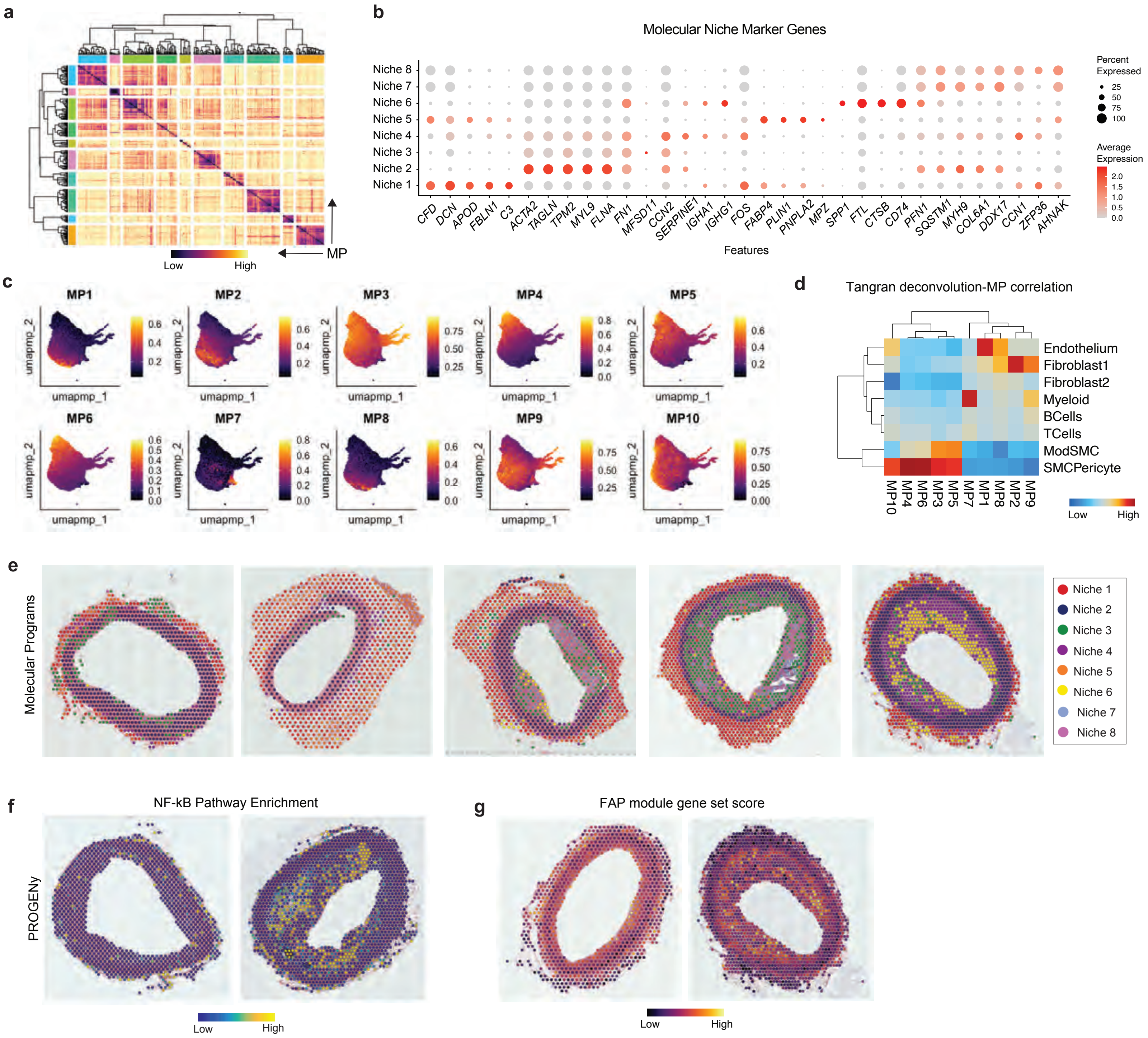

### Supplemental Figure 7

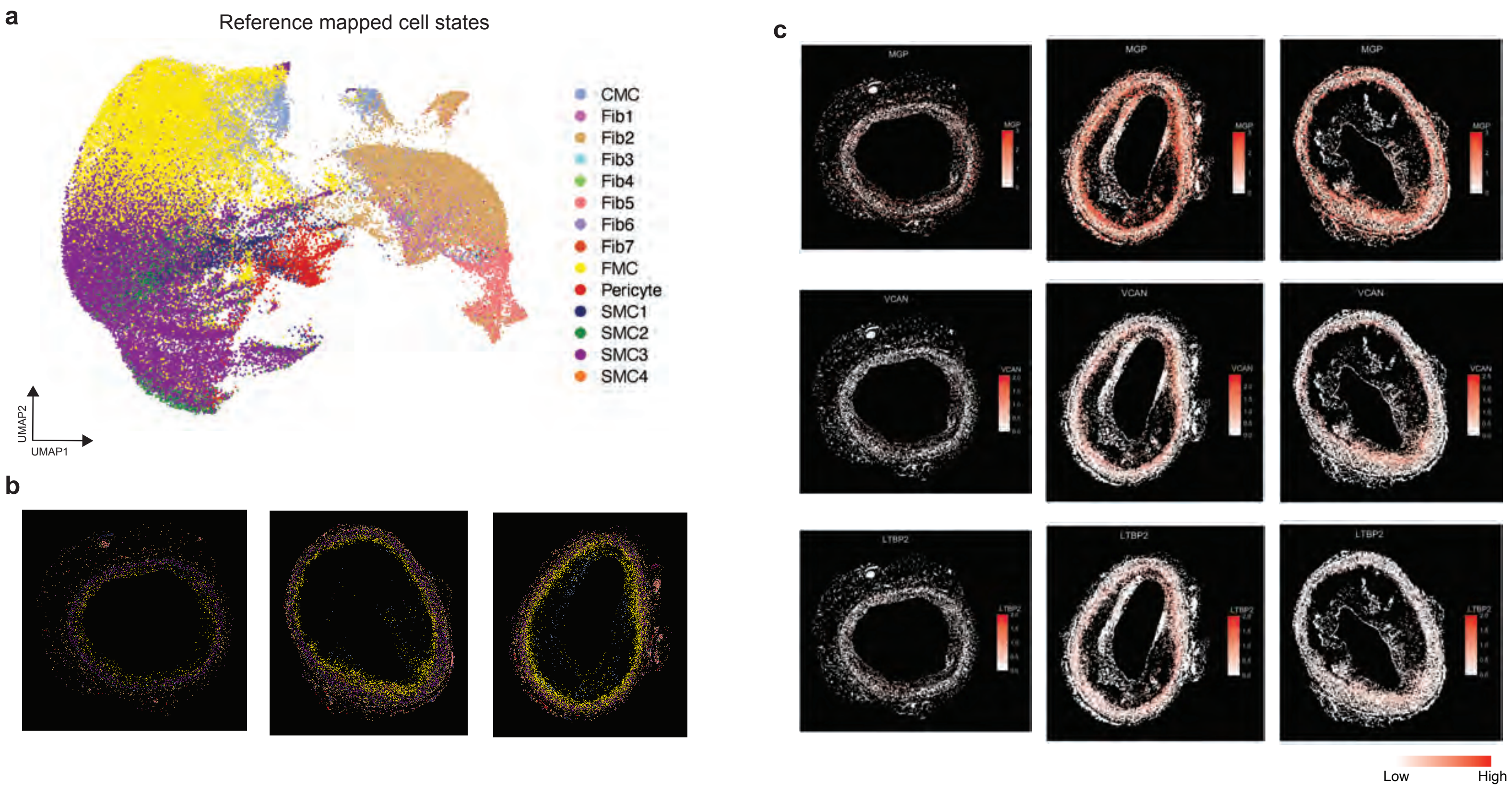

### Supplemental Figure 8

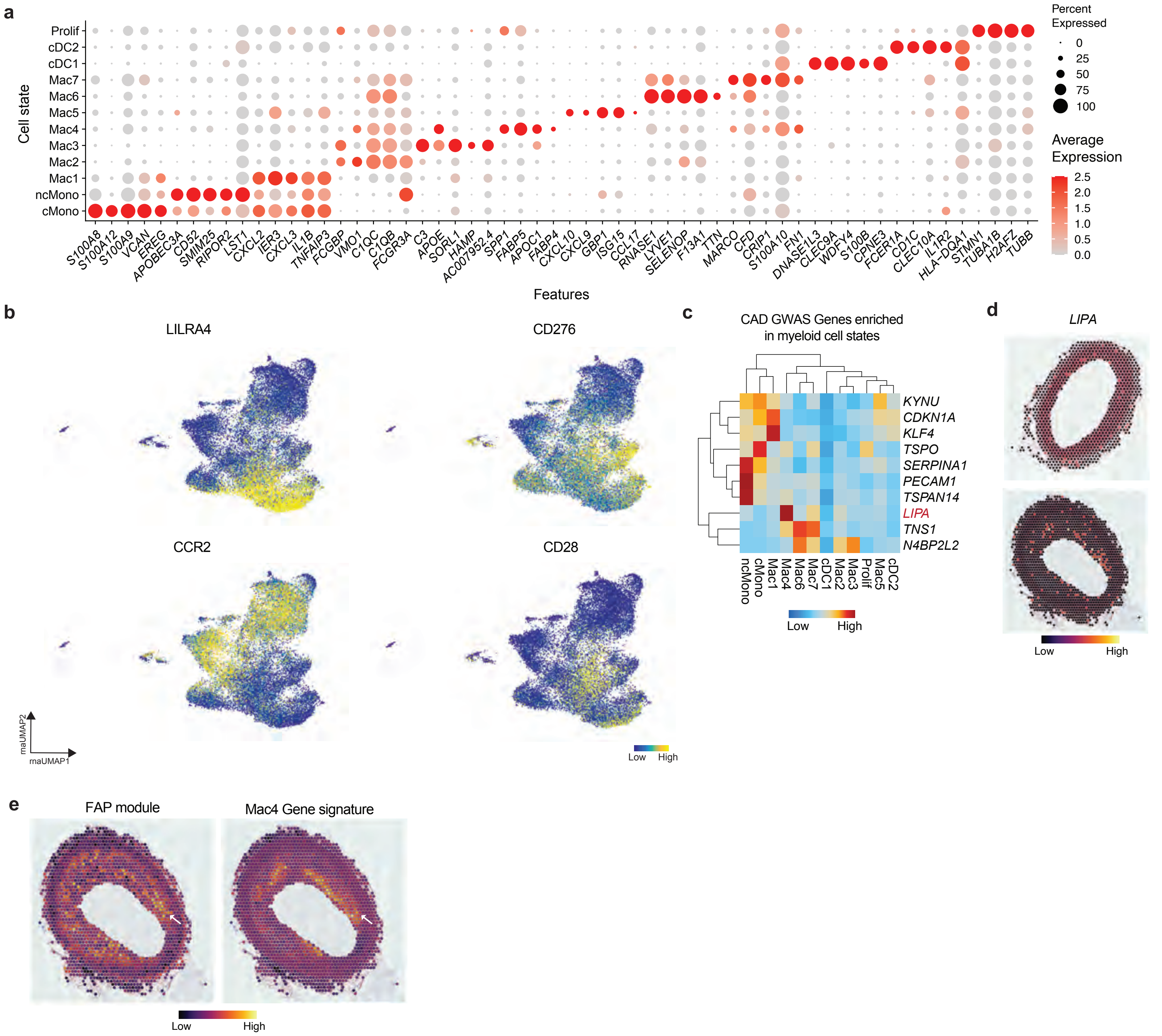

### Supplemental Figure 9

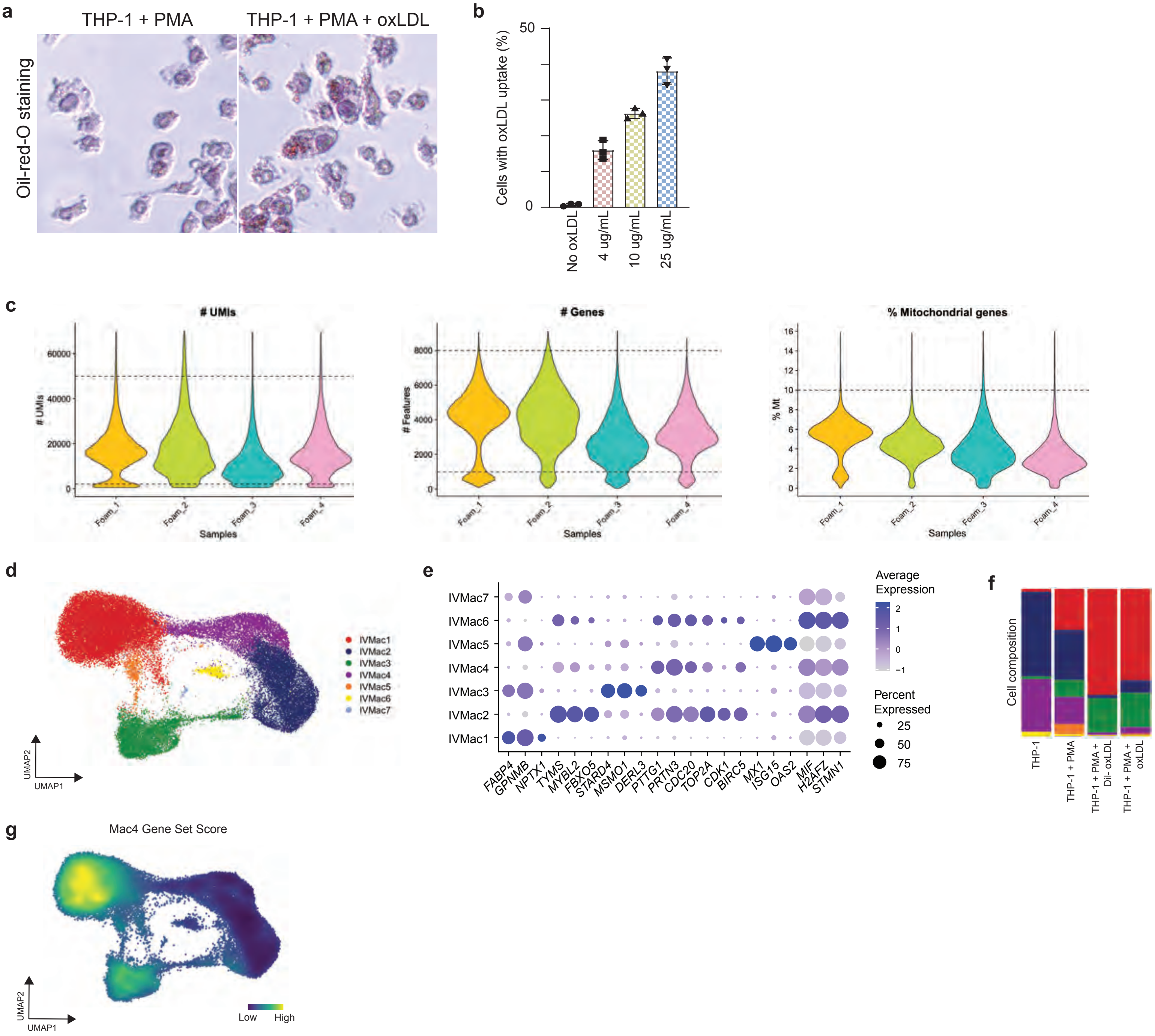
